# Genetic analysis of early phenology in lentil identifies distinct loci controlling component traits

**DOI:** 10.1101/2021.08.16.456438

**Authors:** Vinodan Rajandran, Raul Ortega, Jacqueline K. Vander Schoor, Jakob B. Butler, Jules S. Freeman, Valerie F.G. Hecht, Willie Erskine, Ian C. Murfet, Kirstin E. Bett, James L. Weller

## Abstract

Reproductive phenology is well known to be a key feature of crop adaptation to diverse ecogeographic variation and management practices. Lentil is one of the founder pulse crops of middle-eastern Neolithic agriculture, and the modern-day domesticated lentil germplasm is generally considered to form three broad adaptation groups: Mediterranean, South Asian and northern temperate, which correspond approximately to the major global production environments. Understanding the molecular basis of these adaptations is crucial to maximise efficiency of breeding programs. Here, we use a QTL approach to dissect the earliness that is characteristic of the South Asian *pilosae* ecotype, and that suits it to the typically short winter cropping season. We identified two loci, *DTF6a* and *DTF6b*, at which dominant alleles confer early flowering. We show that, although these loci can interact in an additive manner, *DTF6a* alone is sufficient to confer early flowering even in extremely short photoperiods. Comparisons with closely related legume species confirmed the presence of a conserved cluster of three *FT* orthologs among potential candidate genes in the region, and expression analysis in near-isogenic material showed that the early *dtf6a* allele is associated with a strong derepression of the *FTa1* gene in particular. Analysis of sequence variation revealed the presence of a 7.4 kb deletion in the *FTa1*-*FTa2* intergenic region in the *pilosae* parent, and a wide survey of over 400 accessions with diverse origin showed that the *dtf6a* allele is dominant in South Asia material. Collectively, these results contribute to understanding the molecular basis of global adaptation in lentil, and further emphasize the importance of this conserved genomic region for adaptation in temperate legumes generally.

## INTRODUCTION

Lentil (*Lens culinaris*) is a grain legume that is widely cultivated around the world and is a staple source of dietary protein and fibre in many countries (Iqbal et al. 2006). Global production is dominated by Canada and India, but substantial production also occurs in Australia, Turkey and Nepal (FAO 2020). As in many crop species, appropriate phenology in lentil is a critical determinant of yield and is an important consideration in the development of varieties suited to particular production environments (Hodges 1990; Kumar et al 2016).

Lentil was domesticated in the Fertile Crescent from the wild species *L. orientalis*, a vernalization-responsive long-day species with a winter annual habit. This phenology was retained in domesticated lentil, suiting it to an autumn sowing cycle. Its expansion west around the Mediterranean and east into central Asia is likely to have required no major change in phenology. However, its subsequent spread south to the Indian subcontinent and Ethiopia, and north to the Caucasus and beyond, likely depended on significant phenological changes suited to different seasonal climates and cropping patterns.

Some insight into this variation was provided by physiological studies which distinguished independent influences of photoperiod and temperature across diverse lentil accessions and described flowering behaviour with a simple photo-thermal model (Summerfield et al. 1985; Erskine et al. 1990; Wright et al. 2021). These studies also observed variation in relative responsiveness to photoperiod and temperature, as well as in time-to-flower under the most inductive conditions, identifying several lines showing minimal acceleration of flowering with increases to temperature, and several others relatively unresponsive to photoperiod (Erskine et al. 1990). A third factor broadly important in the phenology of many temperate annual species, vernalization requirement, is relatively unexplored in lentil, and it is currently not clear how it may relate to photoperiod and temperature responsiveness (Summerfield et al. 1985).

There are currently considered to be three broad adaptation groups of domesticated lentil germplasm; Mediterranean, South Asian and northern temperate groups, which correspond approximately to the major ecogeographic environments in global production. Adaptation to these environments is associated with differences in phenology that reflect differences in relative sensitivity to photoperiod and temperature (Erskine et al. 1994). For instance, when lentil landraces originating from West Asia are grown in Pakistan and India they flower later than local material, which seems to be generally less sensitive to photoperiod and more sensitive to temperature (Erskine and Saxena 1993; Erskine et al. 1994). This reduced sensitivity to photoperiod in South Asian landraces is suggested to improve adaptation at lower latitudes by ensuring that flowering occurs at shorter daylength, thus minimizing risk of exposure to late season drought and accommodating the crop efficiently within an intense production system (Erskine et al. 1994).

Such differences in phenology remain a significant barrier to the use of exotic material for germplasm improvement, and a better understanding of their genetic basis and environmental drivers is of high importance for breeding. For example, the narrow genetic base of South Asian germplasm has been substantially broadened by the introduction of elite higher-yielding lines that carry an exotic source of photoperiod-insensitive flowering and were thereby pre-adapted to the local environment (Erskine et al. 1998; Sarker et al. 1999). Another example can be seen in the integration of climate data, the photothermal model and knowledge of flowering time diversity to facilitate selection of genotypes suited to winter sowing in the highlands of central and eastern Turkey (Keatinge et al. 1995; Keatinge et al. 1996).

In the South Asian case, the novel genetics behind the alternative adaptation was shown to derive primarily from a recessive allele at the *Sn* locus (Sarker et al. 1999), which was subsequently identified as the lentil ortholog of the Arabidopsis circadian clock gene *ELF3* (Weller et al. 2012). However, little is known about the genetic basis for the adaptation of the indigenous South Asian material (also referred to as *pilosae* ecotype). In the present study, we first aimed to characterize the variation in flowering response to photoperiod and vernalization of sixteen lentil accessions representing the full range of variation in photoperiod response (Erskine et al. 1990). We then sought to define the genetic basis for the distinct early flowering of a representative South Asian accession.

## METHODS

### Plant material and growth conditions

Fifteen accessions of *Lens culinaris* Medik. and one accession of its putative wild progenitor *L. orientalis* (Boiss.) Handel-Mazetti were obtained from the collection at the International Center for Agricultural Research in Dry Areas (ICARDA) (**Supp Table 1**). For each accession, 24 seeds were scarified and sown two per pot in 140mm pots, in a 1:1 mix of vermiculite and dolerite chips topped with approx. 3cm of sterile nursery grade potting mix with controlled release fertilizer and granulated sand. Half of the seeds were subjected to a 32-day vernalization treatment at 4°C, applied to imbibed seed. Unvernalized seeds were sown several days prior to the end of this period in order to synchronize emergence in the two treatment groups. Material was then grown in short (12-h) or long (16-h) day conditions, with six plants subjected to each factorial combination of photoperiod and vernalization treatments. All plants received 8-h of natural daylight in a heated glasshouse (mean day temperature 23°C) before automated daily transfer to night compartments held at 16°C in which they received a 4-h photoperiod extension with white light from a combination of fluorescent tubes (50µmol m^-2^s^-1^) and incandescent globes (5µmol m^-2^s^-1^). This extension was followed either by 12-h darkness (SD) or a further 4-h extension with light from the incandescent globes alone (LD). Plants were lightly watered regularly, and a nutrient solution applied weekly.

For genetic mapping and association analysis, an F_2_ population (n=173) was generated from the cross between the early-flowering Indian landrace ILL 2601 and the late-flowering accession ILL 5588 (cv. Northfield). This population was sown in February 2013 (with growing medium and maintenance as described above) at the University of Tasmania phytotron and evaluated under a short-day photoperiod as described above, while parental accessions were in addition evaluated under a long day photoperiod. A total of nine traits related to phenology, plant architecture and seedling emergence were evaluated during the growing season for use in QTL analysis.

Three pairs of F_4_ near-isogenic lines (NILs) segregating for either *DTF6a* (NIL9 and NIL52) or *DTF6b* (NIL85) were developed from progeny of single F_2_ individuals derived from the cross ILL 2601 x ILL 5588, by marker-assisted selection of appropriate recombinants in subsequent generations.

### Genotyping, linkage map construction and QTL analysis

Genotyping was performed by Diversity Array Technology Pty. Ltd. (Canberra, Australia) using DArTseq technology (Sansaloni et al. 2011), which generates 64 base pairs (bp) of sequence at each marker by next generation sequencing. The 173 individuals of the ILL 2601 x ILL 5588 F_2_ mapping population were genotyped with a total of 9,315 markers. Markers that were identified to be heterozygous for either parent, non-polymorphic, or with > 2% missing data were excluded from subsequent analysis. A subset of 2,161 polymorphic markers were retained and utilised in the construction of the linkage map.

A genetic linkage map was constructed for the mapping population using JoinMap 4.0 (Van Ooijen 2006), using markers without significant segregation distortion and < 95% similarity to any other marker. A minimum logarithm of odds (LOD) value of 10.0 was used as the significance threshold to assign markers to groups. The regression algorithm was applied with the Kosambi mapping function and default settings to estimate the order of the polymorphic markers and the distances between markers within each group. After the first iteration of regression mapping, markers within ±1cM of another marker were manually removed (to reduce marker numbers in high density regions) and another iteration of regression mapping was employed. Markers with a nearest neighbour fit of > 6cM or a genotype probability < 5.0 (-log_10_(P)) were progressively excluded from each linkage group. In linkage groups with a larger number of excluded markers (> 20% of total markers), a secondary attempt at map construction was undertaken including markers with segregation distortion and high similarity. The nomenclature proposed for the linkage groups is adapted from Sharpe et al. (2013). The linkage map was visualised using MapChart 2.3 (Voorrips 2002). To ascertain the syntenic relationship of the lentil linkage groups obtained in this study with that of *M. truncatula*, the sequences of the DArT-Seq markers incorporated into the final framework of the ILL 2601 x ILL 5588 genetic linkage map were used in a BLAST search against the Medicago reference genome version Mt4.0 (Tang et al. 2014).

QTL analysis on the traits scored in the F_2_ population was undertaken using MapQTL 6 (Van Ooijen, 2009). Permutation tests were run to determine LOD significance thresholds at genome-wide levels (1,000 permutations) (Churchill and Doerge 1994). QTL analyses were first performed using interval mapping. For each putative QTL exceeding the significance threshold in interval mapping, the marker closest to the QTL peak was chosen as a cofactor for multiple-QTL model (MQM) mapping. Cofactors were initially user-determined and then subject to a likelihood analysis based on backward elimination (*p* < 0.05) employed by the automatic cofactor selection function to determine their suitability for MQM analysis. The MQM analyses were performed iteratively until no new QTL were detected and QTL positions were stable (Van Ooijen 2009). The effect of the peak marker at each QTL on the associated phenotype was also examined.

### Gene expression studies

For the expression analysis, two pairs of NILs segregating at the single locus *DTF6a* were grown under short day (12-hr) photoperiod in an automated phytotron at the University of Tasmania. For quantitative reverse transcriptase PCR (qRT-PCR), samples of the most recent fully expanded leaves were harvested three weeks after emergence. RNA extraction, cDNA synthesis and gene expression determination were performed as described in Sussmilch et al. (2015) using the primers indicated in **Supp Table 2**. The expression level of tested genes was normalized against *Actin* using the ΔΔCt method.

## RESULTS

### Variation in lentil responsiveness to photoperiod and vernalization

We initially surveyed the variation in flowering time (DTF, the number of days to the opening of the first fully developed flower) and responses to photoperiod and vernalization in a small but diverse collection of lentil accessions representative of major adaptation groups (**Supp Table 1**). **Figure 1A** shows that these accessions displayed a diverse range of flowering times, ranging from 34 to 50 d under LD and from 43 to 145 d under SD. Interestingly, flowering times under these two conditions were not correlated (**Figure 1B**), suggesting they are likely subject to independent genetic control. However, flowering time under SD did show a strong correlation (r^2^=0.98) with photoperiod response for DTF (i.e. the difference between DTF in SD and in LD), which also varied widely, from 1.4 d in ILL 6005 to over 100 d in ILL131 and ILL 1756 (**Figure 1C**). Among the 16 accessions, three (ILL 4605, ILL 6005 and ILL 2601) were distinctively early-flowering and showed near-insensitivity to photoperiod (**Figure 1C**). A second group of five accessions, including the wild *L. orientalis* line ILWL 7, were late flowering with a strong photoperiod response, and the remaining eight accessions showed intermediate responsiveness.

**Figure 1.**
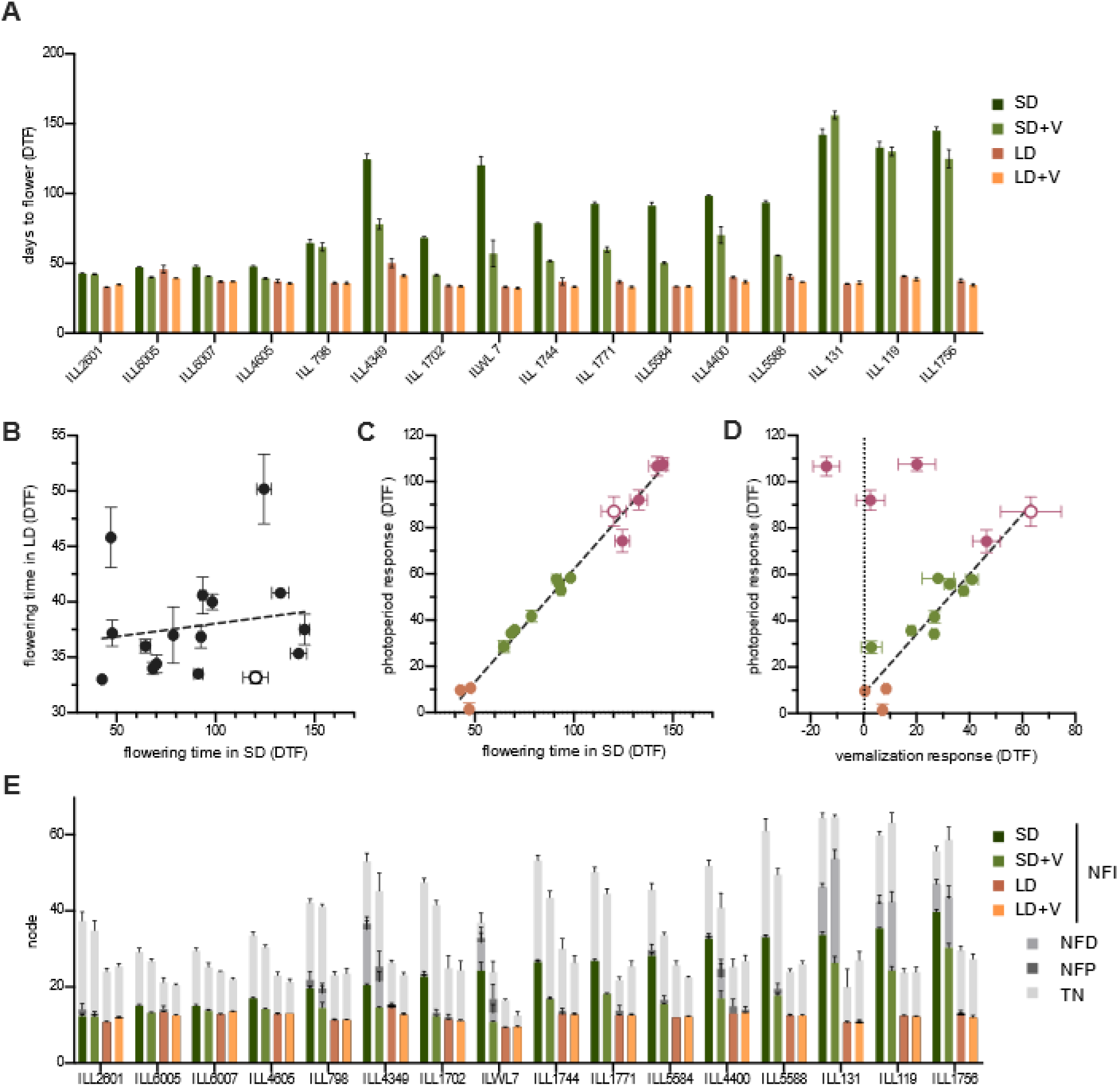
Variation in flowering time and other aspects of reproductive development in response to photoperiod and vernalization. **(A)** Flowering time of lentil accessions under short (SD) or long day (LD) conditions, either with (+V) or without a vernalization treatment. **(B)** Relationship between flowering time in SD and LD. **(C)** Relationship between photoperiod response and flowering time in SD. Possibly distinct groupings are indicated in orange (early-flowering, low sensitivity), maroon (late-flowering, high sensitivity) and green (intermediate sensitivity). **(D)** Relationship between photoperiod response and vernalization response for DTF. Colours represent the same groupings as in (C). **(E)** Phasing of reproductive development. Coloured bars represent the node of flower initiation (NFI). Stacked bars in shades of grey represent the node of first developed flower (NFD), node of first pod (NFP) and the total number of nodes (TN) at maturity (terminal arrest). The wild *L. orientalis* accession ILWL 7 is indicated by open symbols in panels A-C. All values represent mean ± SE for n=4-6.

We also examined the response of this panel to a standard vernalization treatment in which imbibed seed were maintained at 4°C for four weeks. The strongest response, a >60 d promotion of flowering, was seen in ILWL 7, and the majority of accessions showed a tight relationship (r^2^=0.87) between their degree of responsiveness to photoperiod and vernalization for DTF (**Figure 1D**). However, the three latest-flowering accessions under SD (ILL 119, ILL 131 and ILL 1756) showed a disproportionately weak vernalization response, potentially identifying them as a distinct response class. Of these, one (ILL 119) had no significant vernalization response, and across all remaining accessions, two others (ILL 4605 and ILL 798) also showed no significant response (**Figure 1D**).

In addition to the simple DTF measurement, we also recorded other aspects of reproductive development in these accessions, including the node of initiation of the first floral structure, (whether a fully-developed flower or an arrested/aborted bud) (NFI), the node at which the first flower developed (NFD), the node at which the first pod was produced (NFP) and the total number of nodes at apical arrest (TN). These measurements allowed us to quantify the developmental intervals over which initiated flower buds failed to progress to formation of open flowers and to pods, and the duration of the flowering phase. Flower bud abortion is anecdotally important in several crop legumes (Sita et al. 2017; Maqbool et al. 2010), and in pea has been a characteristic and diagnostic feature of certain allelic combinations for established flowering time loci (Murfet, 1985).

**Figure 1E** shows that under SD conditions, several lines showed significant early abortion of flower buds and/or fully-formed flowers. This tendency was most prominent in the accessions that were latest-flowering in SD and showed the strongest photoperiod response for DTF (**Figure 1B**), suggesting that these two responses could be a manifestation of a single underlying physiological or genetic system controlling initiation and development.

### Genetic analysis of early flowering in ILL 2601

The reduced photoperiod sensitivity of cv. Precoz (ILL 4605) and derivatives such as ILL 6005 was initially shown to be conferred by recessive alleles at the *Sn* locus (Sarker et al. 1999). In an earlier study, we confirmed this finding in progeny of a cross between ILL 5588 (cv. Northfield) and ILL 6005, and identified a co-segregating mutation in an *ELF3* ortholog as the likely cause of the photoperiod-insensitive phenotype (Weller et al. 2012). Among the accessions examined in **Figure 1**, only line ILL 2601 showed a similar dramatic loss of photoperiod sensitivity to Precoz and ILL 6005. In contrast to the progeny of a cross between ILL 4605 and ILL 6005, the F_2_ of a cross between ILL 2601 and ILL 6005 segregated late-flowering individuals under short days (J. Weller, unpubl.) indicating a genetic basis for earliness independent of *Sn*. This conclusion was further supported by sequencing of the *ELF3a* gene from ILL 2601, which revealed no functionally-significant differences relative to ILL 5588 (**Supp Figure 1**).

To characterize the genetic control of the earliness in ILL 2601, we crossed it to ILL 5588 and assessed an F_2_ population of 173 individuals under 12-h SD conditions. In addition to the traits described above, we also recorded time to seedling emergence (days to emergence; DTE) given that variation for this trait had been noted in some circumstances previously and could potentially complicate a flowering time assessment based on sowing date alone. Subsequently, we recorded DTF as starting from the date of seedling emergence. We also measured total plant height (PH), number of branches >5 mm in length at 3 weeks post-emergence (number of early branches; EBN), total length of these branches (early branch length; EBL) and the distance between nodes 1 and 9 on the main stem (INL).

In the F_2_ population, DTF showed an essentially continuous distribution (**Figure 2A**), although skewed towards the early phenotype of ILL 2601. This suggests the presence in ILL 2601 of at least one dominant allele conferring early flowering. In addition, the distribution showed a degree of transgressive segregation relative to the parental range in both directions, suggesting the presence of other minor loci potentially involved in the control of DTF. A strong correlation (R^2^=0.905) was found between DTF and NFD, but the correlations between DTF and NFI (R^2^=0.407) and NFI and NFD (R^2^=0.4875) were weaker, reflecting the occurrence of early flower abortion in several individuals (**Supp Figure 2**).

**Fig. 2.**
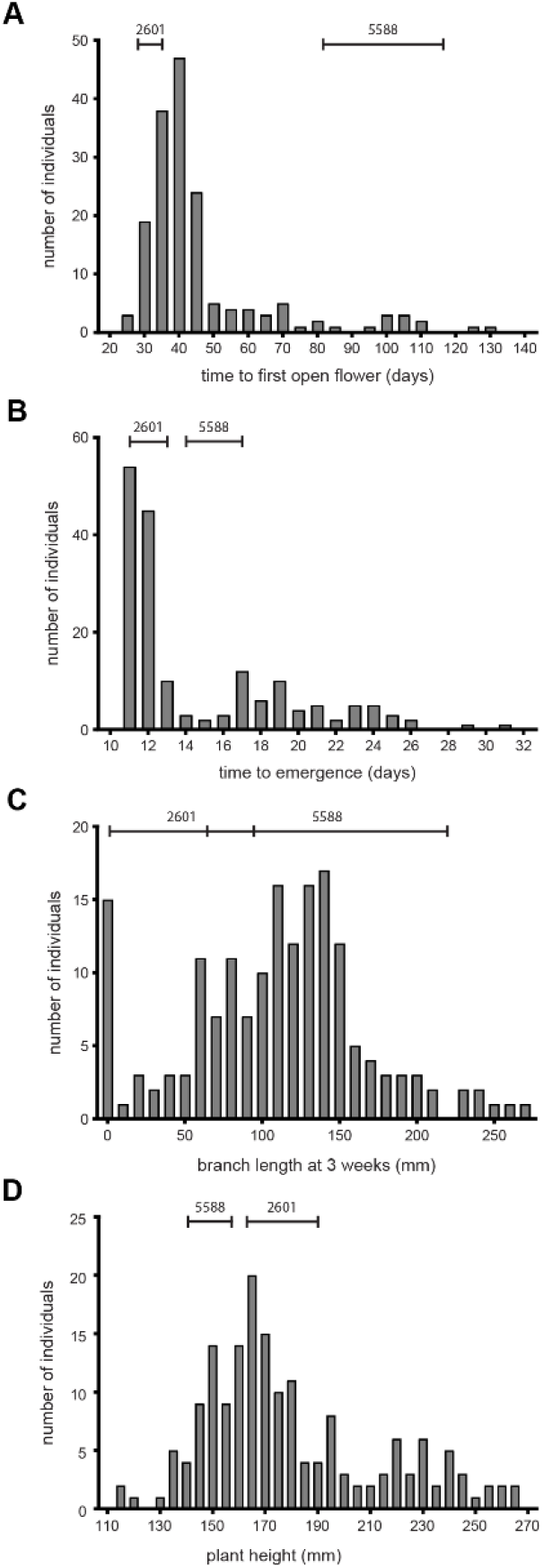
Frequency distributions for phenology and growth habit traits in an F_2_ population. Days to flower **(A)**, days from sowing to seedling emergence **(B)**, total branch length at three weeks post-emergence **(C)** and total plant height **(D)** were recorded in the F_2_ progeny of a cross between accessions ILL 2601 and ILL 5588. The ranges in parental values are indicated by horizontal bars.

Analysis of other traits DTE, EBL and PH in the F_2_ generation showed a continuous distribution (**Figure 2 B, C, D**) confirming these as quantitative characters, although there was an indication of bimodality for DTE and PH, implying the involvement of major loci. For both DTE and PH, the range of values in the population was much wider than that in the parental lines, indicating a recombination of parental alleles at multiple loci. For DTE, only late transgressive segregants were recorded, indicating that the allelic combination present in ILL 2601 specified the minimal emergence time, but also included alleles capable of contributing to delayed emergence.

### Linkage mapping and QTL analysis

To further investigate the genetic control of the observed variations, we constructed a genetic linkage map for the ILL 2601 x ILL 5588 F_2_ population using a total of 734 DArT-seq markers (**Supp Table 3, Supp Figure 3**). The final map has an overall length of 1032 cM, defined by seven linkage groups that correspond to the seven chromosomes in the lentil genome (2n = 14). The average distance between adjacent markers was 1.41 cM, with only one interval greater than 10 cM (**Supp Table 4**).

QTL analysis on flowering traits yielded a total of nine QTLs distributed on LG6 and LG2 (**Table 1**); Of these, seven out the nine were detected on LG6. The node of flower initiation (NFI) was associated with only one locus, in a central region of LG6 (LG6A). This region also influenced days to flowering (DTF) along with a second distinct region on LG6 (LG6B) (**Figure 3**). Together with a third region on LG2, both LG6A and LG6B regions also contributed to control the node of flower development (NFD) and the initiation-development interval (DFD). In general the effects of the QTLs detected on LG6 were stronger than those detected on LG2 for each trait, with estimates of observed variation explained as high as 46% (DTF) or 49% (NFD) when QTLs for these traits in LG6 are combined. In comparison, the LOD and PVE values obtained for the QTLs in LG2 were lower, indicating a contribution of only around 10%.

**Table 1.** Quantitative trait loci for flowering time traits detected in the ILL2601 x ILL5588 progeny.

| **Trait** | **QTL** | **LG** | **QTL peak position (cM)** | **LOD** | **Variation explained (%)** | **Peak marker** | **Marker position (cM)** |
| --- | --- | --- | --- | --- | --- | --- | --- |
| Days to flowering (DTF) | *qDTF6a* | 6 | 67.3 | 17.6 | 33.9 | 101290509 | 68.6 |
|  | *qDTF6b* | 6 | 152.3 | 7.1 | 12.0 | 3659911 | 152.8 |
| Node of flower development (NFD) | *qNFD6a* | 6 | 67.3 | 13.1 | 23.4 | 101290509 | 68.6 |
|  | *qNFD6b* | 6 | 152.3 | 8.7 | 15.0 | 3659911 | 152.8 |
|  | *qNFD2* | 2 | 68.1 | 6.4 | 10.5 | 3632005 | 68.1 |
| Delay to flower development (DFD) | *qDFD6a* | 6 | 67.3 | 6.3 | 12.6 | 101290509 | 68.6 |
|  | *qDFD6b* | 6 | 152.3 | 7.8 | 16.1 | 3659911 | 152.8 |
|  | *qDFD2* | 2 | 68.1 | 4.2 | 8.2 | 3632005 | 68.1 |
| Node of floral initiation (NFI) | *qNFI6a* | 6 | 67.3 | 8.0 | 20.0 | 101290509 | 68.6 |

**Fig. 3.**
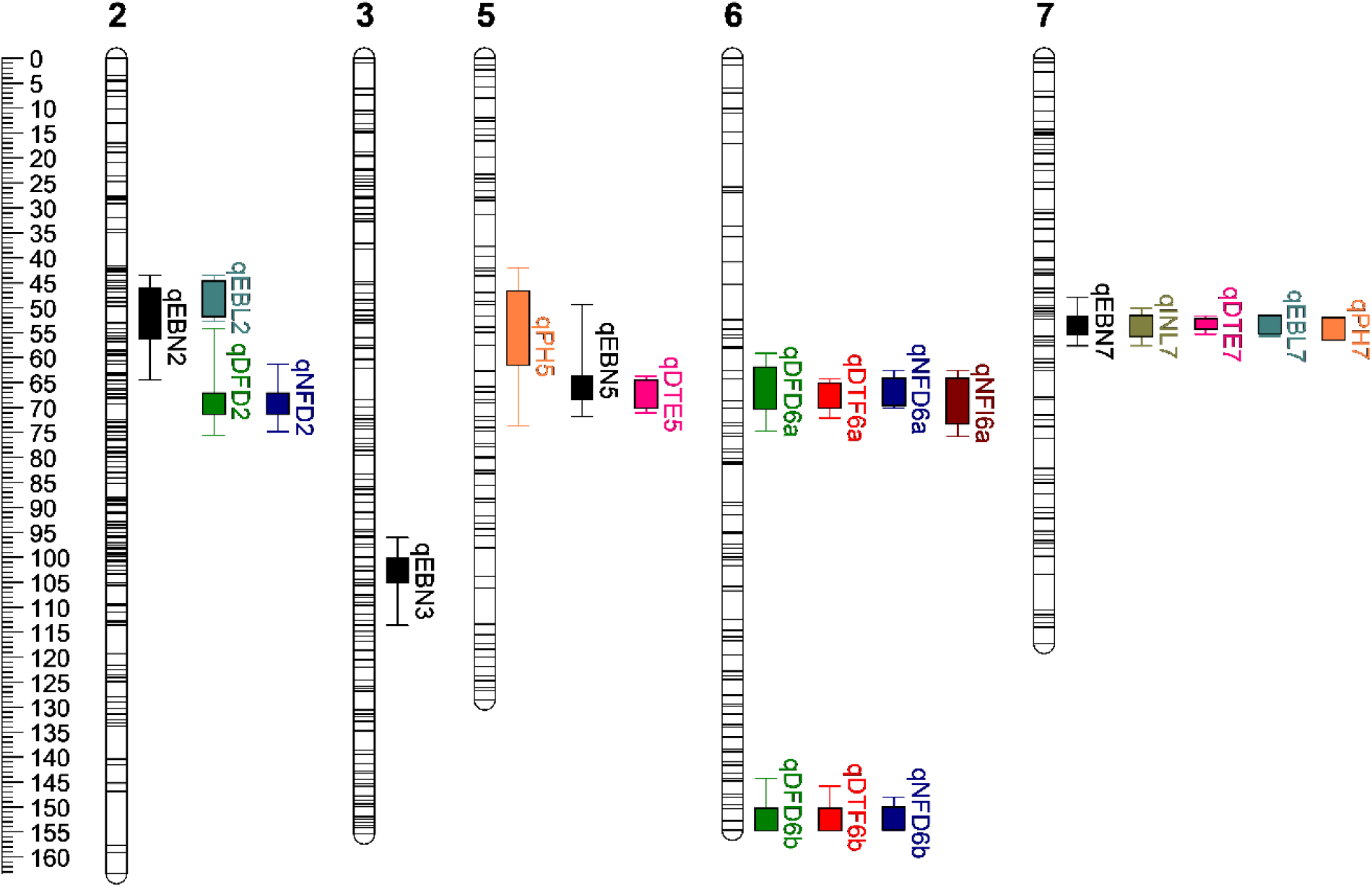
Linkage map showing QTL detected in the ILL 2601 x ILL 5588 population. Scale is cM. QTL nomenclature follows Tables 1 and 2. Box and whiskers represent 1-LOD and 2-LOD intervals, respectively, around each QTL peak.

Three of the four QTL in the LG6A region (*qDTF6a, qNFD6a* and *qNFI6a*) are the largest contributors to their respective traits, estimated to explain over 20% of the observed variation in each case (**Table 1**). QTL in the second LG6B region had mostly smaller but still substantial effects, explaining an estimated 12-16% of the observed variation. In both regions, QTLs were observed to either occur at the same position (peak marker) or were co-located within ±1 cM (**Figure 3**) and, given the physiologically related nature of the traits concerned, it seems most likely that they represent a single underlying molecular cause in each region. Although this conclusion must be proven empirically, for convenience we will provisionally refer to these two loci as *DTF6a* and *DTF6b*.

Eleven QTLs were detected for traits related to seedling emergence and plant architecture (**Table 2**); two each for plant height (PH), emergence (DTE) and the length of early branches (EBL), four for the number of early branches (EBN), and one for length between nodes 1 and 9 on the main stem (INL). Of these QTLs, five were located on LG7 in the same interval. In this region, the QTL *qDTE7, qPH7* and *qINL7* each explained >20% of the observed variation.

**Table 2.** Quantitative trait loci for other traits detected in the ILL2601 x ILL5588 progeny.

| **Trait** | **QTL** | **LG** | **QTL peak position (cM)** | **LOD** | **Variation explained (%)** | **Peak marker** | **Marker position (cM)** |
| --- | --- | --- | --- | --- | --- | --- | --- |
| Days to emergence (DTE) | *qDTE7* | 7 | 53.3 | 35.4 | 53.7 | 3631532 | 52.3 |
|  | *qDTE5* | 5 | 68.5 | 11.5 | 12.6 | 3635263 | 68.5 |
| Number of early branches (EBN) | *qEBN7* | 7 | 53.3 | 7.9 | 14.5 | 3631532 | 52.3 |
|  | *qEBN5* | 5 | 65.8 | 5.4 | 9.6 | 3634578 | 65.8 |
|  | *qEBN3* | 3 | 101.8 | 4.9 | 8.6 | 3630041 | 101.8 |
|  | *qEBN2* | 2 | 49.7 | 4.7 | 8.3 | 3634748 | 49.7 |
| Total length of early branches (EBL) | *qEBL2* | 2 | 47.8 | 6.5 | 14.3 | 3630240 | 46.8 |
|  | *qEBL7* | 7 | 52.3 | 5.2 | 11.1 | 3631532 | 52.3 |
| Plant height (PH) | *qPH7* | 7 | 55.3 | 14.8 | 30.6 | 3630555 | 55.8 |
|  | *qPH5* | 5 | 52.7 | 4.4 | 7.9 | 3634509 | 53.9 |
| Length between nodes 1 and 9 on the main stem (INL) | *qINL7* | 7 | 53.3 | 8.4 | 20.4 | 3631532 | 52.3 |

### Interaction between *DTF6a* and *DTF6b*

To better understand the nature and functional relationship of *DTF6a* and *DTF6b* in control of flowering time, we next examined their individual allelic effects and their interaction in the F_2_ population, as inferred from peak marker genotypes. In view of the strong photoperiod response of the wild *L. orientalis* and of *L. culinaris* accessions from the wider domestication region (**Figure 1, Supp Table 1**), the alleles from ILL 5588 can reasonably be considered ancestral, and for the purpose of clarity will be referred to as *DTF6a* and *DTF6b*, and the presumed derived alleles from ILL 2601 as *dtf6a* and *dtf6b*.

With a *DTF6b* genetic background, F_2_ individuals heterozygous for *DTF6a* did not differ significantly from *dtf6a* homozygous segregants for DTF. Similarly, with a *DTF6a* background, mean DTF of the *DTF6b dtf6b* heterozygous progeny showed no significant difference compared to *dtf6b* homozygous individuals (**Figure 4A**). Taken together, these results indicate a dominant mode of inheritance of the early flowering variants in the SD conditions tested.

**Fig. 4.**
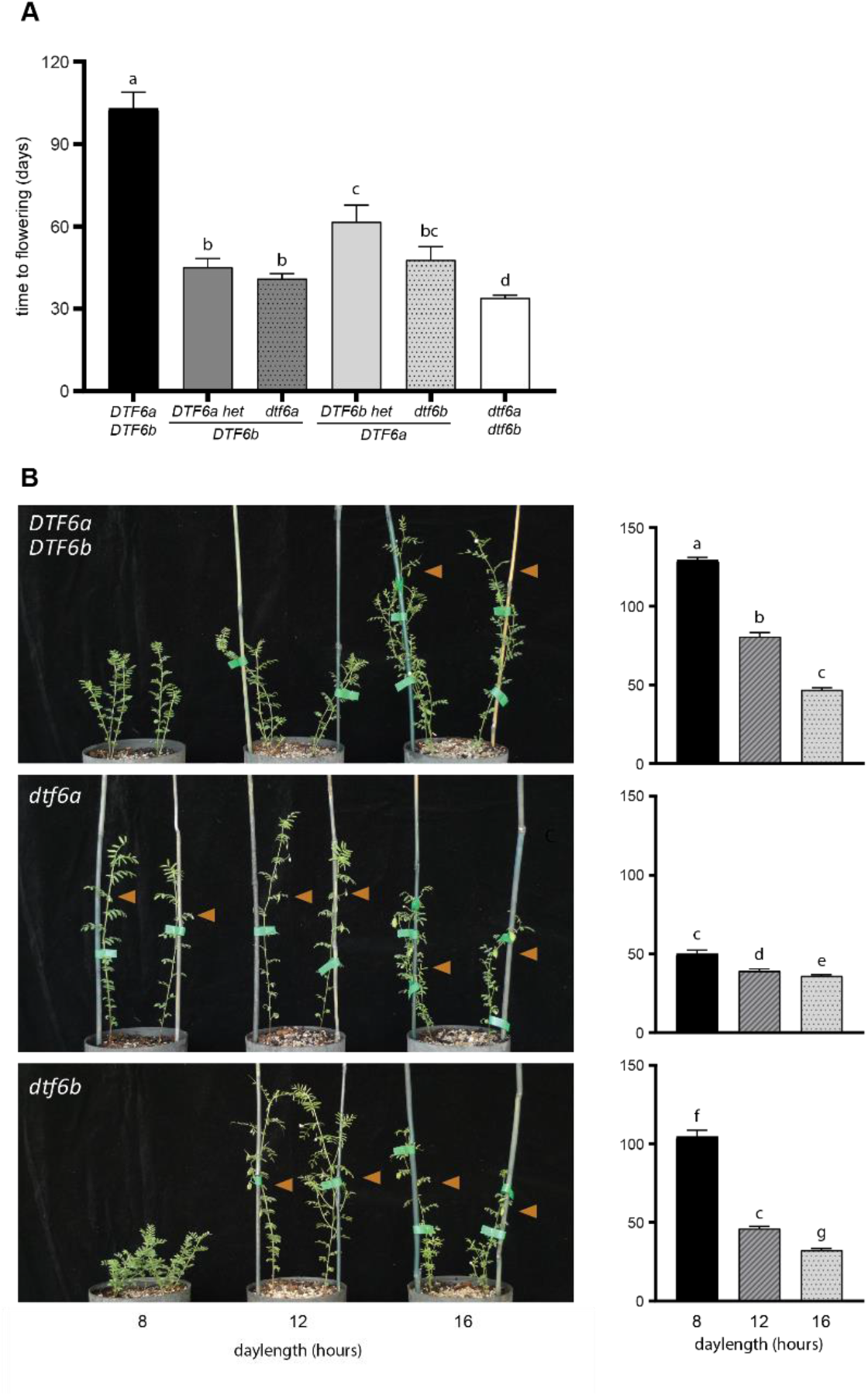
*qDTF6a* and *qDTF6b* peak marker effects under different photoperiods. **(A)** Interaction between *DTF6a* and *DTF6b* for flowering time in the ILL 2601 x ILL 5588 F_2_ population under short day conditions (12h). **(B)** F_4_ NILs differing only at the *DTF6a* (middle panel) or *DTF6b* (bottom panel) locus grown under three different photoperiods: SD (8hr), SD (12hr) and LD (16hr). Plants in the top panel carry ILL 5588 alleles at both loci. Orange triangles indicate the plants that have initiated flowering with flowering time data shown on the right.

The genetic interaction of the two loci was assessed in the F_2_ progeny by comparing mean DTF of the six genotypic classes. The data in **Figure 4A** show that *DTF6a DTF6b* double homozygous progeny flowered significantly later (p < 0.05) than each of the other genotypic classes. While the individual effects of the *dtf6a* and *dtf6b* alleles were indistinguishable (p = 0.345), *dtf6a dtf6b* individuals did flower significantly earlier than those with any other allelic combination (p < 0.05).

For more detailed investigation of the individual effects of *DTF6a* and *DTF6b*, we developed NILs differing either at *DTF6a* or *DTF6b* and examined them under three different photoperiods: 8, 12 or 16h (**Figure 4B**). Under all three photoperiod conditions, the *dtf6a* and *dtf6b* genotypes conferred significantly earlier flowering than *DTF6a DTF6b*, but their effects were much stronger under the two short-day photoperiods (minimum 40d promotion) than under LD (10d earlier), indicating that both loci influenced photoperiod sensitivity. Under 12h, the effects of the individual loci were similar to those seen in the F_2_ population grown under the same conditions. Both *dtf6a* and *dtf6b* genotypes were significantly earlier flowering than *DTF6a DTF6b* plants (p < 0.001 for both), although in this case the *dtf6a* line was slightly earlier than *dtf6b* (p < 0.001). However, under the more extreme short-day of 8h, the d*tf6a* line showed only a minor additional delay in flowering of around 10d, whereas *DTF6a DTF6b* and *dtf6b* plants showed a more substantial delay of >50d. This result suggests that of the two ILL 2601 alleles, *dtf6a* is more effective than *dtf6b* for promotion of flowering under short days, a conclusion also consistent with the greater contribution of *DTF6a* to the variance observed in the F_2_.

### Characterization of candidate genes

Given that *DTF6a* made the strongest contribution to the control of early flowering in ILL 2601, we chose to investigate its identity further, initially by screening the region for potential candidates. To facilitate the selection of candidates within the region surrounding *qDTF6a*, we first established the macrosyntenic relationships between the seven *L. culinaris* LGs defined in the ILL 2601 x ILL 5588 map (**Figure 3**) with that of the eight chromosomes of *M. truncatula* (2n = 16). The genomes were broadly syntenic (**Supp Figure 4**), but showed the previously documented translocations between the ends of the lentil linkage groups 1 and 5 and major inversions in regions of lentil linkage groups 1 and 7 when compared to *M. truncatula* chromosomes 1 and 8 (Gujaria-Verma et al. 2014; Sharpe et al. 2013; Ramsay et al. 2021). These findings are consistent with previous observations and are the most comprehensive analysis of synteny between these two species to date.

This comparison indicated that the *DTF6a* region corresponded to an interval of approximately 4.23 Mbp (627 genes) in the central region of Medicago chromosome 7. This region was also compared with the syntenic regions of chickpea chromosome 3 and pea chromosome 5 (**Figure 5A**), revealing the presence of several genes associated in some way with flowering time control, and in particular, three orthologs of the Arabidopsis florigen (*FT*) gene. A conserved cluster of *FT* genes is located in this syntenic genomic region across the two major crop legume clades; and in pea, Medicago and chickpea the cluster comprises two genes (*FTa1, FTa2)* arranged in tandem and a third (*FTc*) located either adjacent or nearby (**Supp Figure 5**; Hecht et al. 2011; Laurie et al. 2011; Ortega et al. 2019; Weller et al. 2019). A role for *FT* genes in flowering time control is well documented in many plant species, including legumes (Weller and Ortega 2015; Lin et al. 2021) and among the genes in this cluster in the temperate legumes, most evidence points to *FTa1* as being particularly important. (Hecht et al. 2011; Laurie et al. 2011; Ortega et al. 2019).

**Fig. 5.**
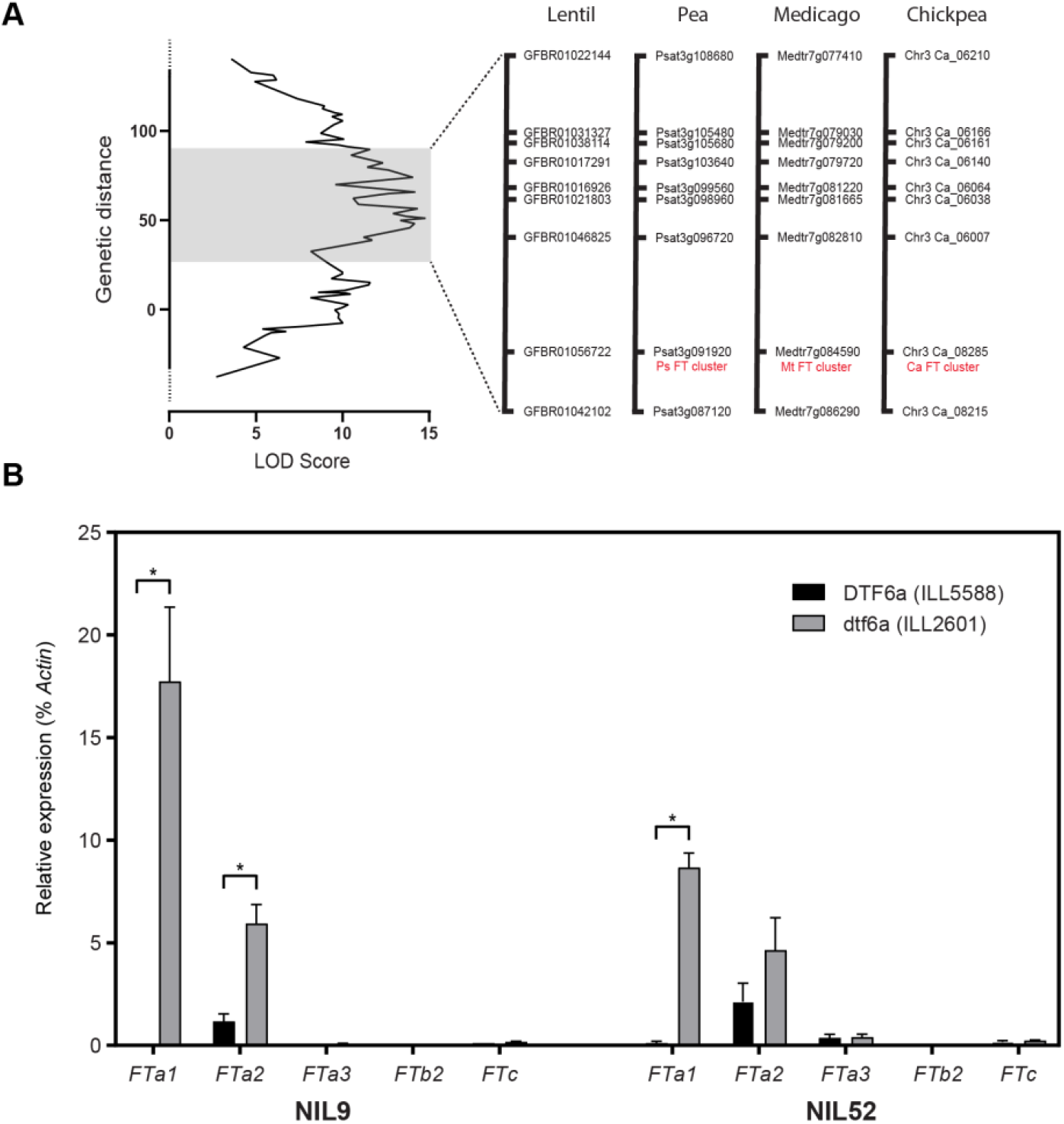
*DTF6a* is located over the conserved *FTa-FTc* cluster and affects its expression. (**A**) QTL peak for *DTF6a*. Lentil markers in the peak region were blasted against Medicago, chickpea and pea to determine the matching genes from the syntenic regions. The *FT* cluster of *FTa1, FTa2* and *FTc* is indicated in red. (**B**) Expression of lentil *FT* orthologues under short day (12hr) photoperiod for two pairs of NILs segregating at the single locus, *DTF6a*. Expression levels show the difference between the photoperiod-sensitive ILL 5588 allele (black) and the early-flowering ILL 2601 allele (grey). The most recent fully expanded leaves were harvested three weeks after emergence. Values have been normalised to the transcript level of *Actin* and represent mean ± SE for n=3 biological replicates, each consisting of pooled material from two plants.

### The *dtf6a* allele is associated with elevated expression of *FTa* genes

Studies of mutants, overexpressors and natural variation have all indicated that *FTa1* genes promote flowering in proportion to their level of expression (Hecht et al. 2011; Laurie et al. 2011; Ortega et al. 2019). Thus, it is possible that the effect of *dtf6a* might be associated with elevated expression or otherwise increased activity of one or more genes in the cluster in the early parent ILL 2601.

To evaluate this possibility, we compared *FT* gene expression in two independently selected pairs of NILs segregating for *DTF6a*, and evaluated their flowering response to photoperiod. As expected, minor differences were observed in DTF of plants grown under LD (**Supp Figure 6**). However, under SD, *DTF6a* plants were unable to flower before the end of the trial, whereas those bearing the *dtf6a* allele flowered at an average of 69.3 (NIL9) and 81.2 (NIL52) days after emergence.

The closely related species pea has six *FT* genes (Hecht et al. 2011; Ortega et al. 2019), and an analysis of recently released lentil genome sequence has confirmed that the same is also true in lentil (Yuan et al. 2021, **Supp Figure 5**). The results in **Figure 5B** show that in leaf tissue of three-weeks old plants grown under SD conditions, both *FTa1* and *FTa2* genes were expressed at a significantly higher level in NILs carrying the *dtf6a* allele compared to equivalent plants with the *DTF6a* allele. *FTa1* expression was undetectable in *DTF6a* NILs and strongly expressed in the *dtf6a* NILs, whereas *FTa2* was expressed at a low level in *DTF6a* lines and only moderately upregulated in one of the two NIL comparisons. In contrast, expression of the four other *FT* genes were not reliably detected above background in any of the NILs. These results imply that the early flowering of *dtf6a* lines is likely to result primarily from an increase in *FTa1* expression, and might reflect the impairment of a cis-acting mechanism normally acting to maintain repression of *FTa1* expression under non-inductive SD photoperiods.

### Sequence variation in the *FTa1*-*FTa2* cluster

To identify sequence variation that might be associated with misregulation of the *FTa1* and *FTa2* genes, we sequenced the *FTa1-FTa2* cluster in the parental lines ILL 2601 and ILL 5588 using an amplicon sequencing approach, from ≈4.5 kb upstream of *FTa1* to ≈1 kb downstream of *FTa2*. The *FTa1* gene is relatively compact, whereas the *FTa2* gene extends to almost 22 kb, with a large third intron over 20 kb in length (21045 bp in ILL 2601, 20723 bp in ILL 5588). We found no sequence differences in the coding sequence of either gene among the two accessions, but identified a total of 136 SNPs and 25 indels distinguishing ILL 5588 and ILL 2601 in non-coding regions (**Supp Figure 7, Supp Tables 7, 8 and 9**). By far the most substantial of these was a large (7,441 bp) deletion that eliminated most of the *FTa1-FTa2* intergenic region in ILL 2601 (**Figure 6 A-C**). Another significant polymorphism, due to its potential position within *FTa1* promoter, is a 245 bp indel found 3,712 bp upstream of the *FTa1* start codon (**Supp Figure 7, Supp Table 7**). Other minor variants were found scattered across the cluster, but they were especially abundant within the third intron of *FTa2*, which accumulated 83% of all SNPs and 56% of all indels detected between the two sequences. This contrasts with the high conservation observed for the *FTa1* gene, where no indels and only 3 SNPs were found within the entire gene (introns included). This contrasting degree of sequence conservation between the two *FTa* genes further support the idea of *FTa1* having a more relevant role than *FTa2* in lentil.

**Fig. 6.**
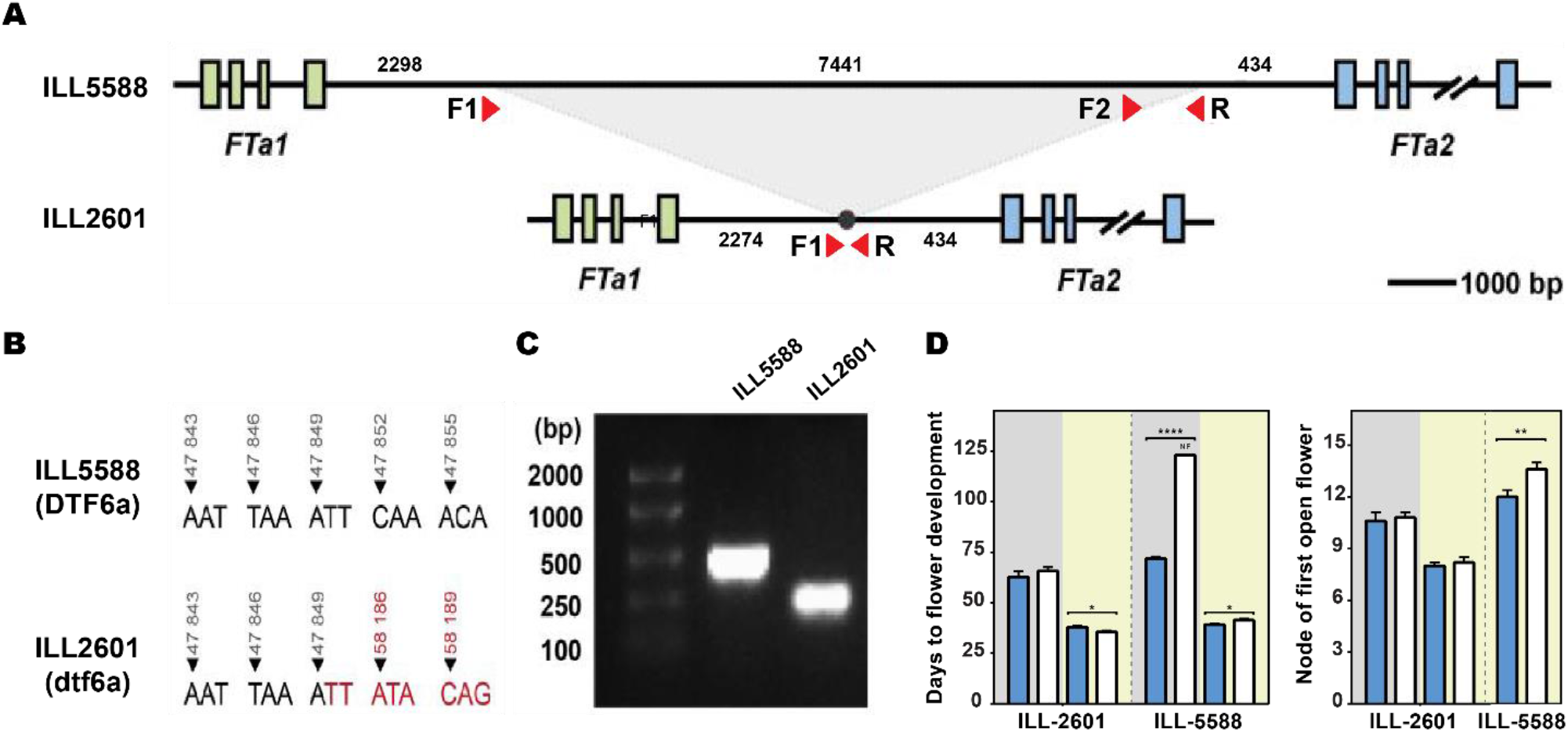
Characterisation of the *FTa1-FTa2* cluster in lentil accessions. **(A)** Schematic diagram of the *FTa1-FTa2* cluster in ILL 5588 and ILL 2601. The green boxes represent the exons of the lentil *FTa1* gene and the blue boxes represent the exons of *FTa2*. **(B)** The sequence position of the 7441-bp deletion in ILL 2601. **(C)** PCR of the intergenic region with a 45s extension time (approximately 1 kb) in ILL 5588 and ILL 2601. Primers positions are annotated in (A). **(D)** Phenotypic response of ILL 5588 and ILL 2601 to photoperiod and vernalization treatment. Blue and white bars represent vernalized and unvernalized plants, respectively. Background colour indicates photoperiod, with SD (8 h) in grey and LD (16 h) in yellow. NF indicates plants unable to flower at the end of the scoring period (123 days). Consequently, their flowering node is not present in the corresponding graph.

In Medicago, transposon insertions in the 3’ region of *FTa1* are reported to confer an early flowering phenotype that is photoperiod sensitive but not responsive to vernalization (Jaudal et al. 2013). To test whether the deletion in the *FTa1*-*FTa2* intergenic region might have a similar differential effect, vernalized and unvernalized plants from ILL 5588 and ILL 2601 were grown under either LD or SD, and flowering time was assessed (**Figure 6D, Supplementary Table 5**). To maximise the differences in flowering time due to photoperiod, a more extreme (8-h) SD treatment was used. In the late-flowering ILL 5588, vernalization caused earlier flowering under both photoperiods. Under SD, unvernalized plants remained vegetative after 123 days, whereas vernalized plants flowered at an average of 71.8 days. Under LD plants flowered earlier and showed a small (2.3 days, 1.3 nodes) but significant (*p* < 0.05) promotion of flowering by vernalization. By contrast, we found no significant reduction in ILL 2601 flowering time due to vernalization, in any of the two photoperiods tested. Moreover, under LD, vernalized plants flowered slightly later than unvernalized plants (2.4 days, *p* < 0.05). However, differences in flowering time and flowering node were notable in response to photoperiod, characterising this line as vernalization-insensitive but photoperiod-responsive.

The physiological similarity between ILL 2601 and the Medicago mutants described above strengthens the idea that the *FTa1*-*FTa2* intergenic deletion might be the causal polymorphism for the observed *FTa1* upregulation. In view of its potential significance, we designed a PCR-based specific marker for this polymorphism (Supplementary Table 10, Supplementary Figure 8), and examined its prevalence in a lentil collection of 48 accessions available at the University of Tasmania and across a broader diversity panel of 324 accessions (Wright et al. 2021), both selected to cover a wide range of geographic origins. Results from the screening were similar in both collections and show that, globally, the deletion is common within lentil germplasm and 23-37.5% of all accessions carry the deletion (**Supplementary Figure 9, Supplementary Figure 10**). However, based on the geographic origin of the accessions, its distribution was uneven: In South Asian material, it was found to be the dominant allelic variant, present in more than 75% of accessions, but it was less frequent in accessions of Middle-Eastern origin and rare in germplasm from Mediterranean and temperate environments (**Supplementary Figure 9, Supplementary Figure 10**). Independently of whether the deletion or any other polymorphism is responsible for the *dtf6a* effect, the high prevalence of the ILL 2601 allele in South Asian accessions suggest that the *DTF6a* locus might substantially contribute to the earliness characteristic of this material.

## DISCUSSION

Identification of variation for target traits and an understanding of the underlying genetics is important for the development of lentil varieties locally adapted to different agro-economical environments. Despite its global importance as a crop, lentil is chronically understudied in this respect. We characterized natural variation for photoperiod and vernalization response in a small collection of lentils representative of the major adaptation groups and found wide variation in the flowering response to these cues (**Figure 1**), in agreement with earlier studies proposing differential photo-thermal sensitivity as a mechanism for latitudinal adaptation (Erskine et al. 1989; Erskine et al. 1990; Erskine et al. 1994). The Indian landrace ILL 2601 was one of the earliest to flower in our survey and has an early phenology typical of the *pilosae* ecotype, which is adapted to the environments of the Indian Subcontinent and is characterized by a reduced photoperiod sensitivity and increased responsiveness to temperature (Erskine et al. 1990; Erskine et al. 1994; Summerfield et al. 1985). Our QTL analysis, performed on a cross between this line and the photoperiod-sensitive accession ILL 5588 (cv. Northfield), detected two loci on lentil chromosome 6 controlling flowering time (*DTF6a* and *DTF6b*). Both loci show a co-dominant-to dominant-early inheritance pattern (**Figure 3, Figure 4A**), and are clearly distinct from the previously described *Sn/ELF3a* locus on chromosome 3. Although both *DTF6a* and *DTF6b* contribute in an additive manner to the phenotype of ILL 2601, analysis of segregants in the F_2_ progeny and near-isogenic lines revealed that the individual effect of the ILL 2601 allele at *DTF6a* is notably stronger than that of the *DTF6b* allele under an 8h photoperiod (**Figure 4B**).

The role of *FT* genes as flowering promoters is widely conserved through the plant kingdom (Wickland and Hanzawa, 2015), and this seems to be true also for legumes (Ortega et al. 2015; Lin et al 2021). In several temperate legumes, flowering time loci show a conserved syntenic position near a cluster of *FT* orthologs, and in chickpea and lupin, are also associated with an increased expression of one or more of the underlying genes (Weller and Ortega, 2015; Nelson et al. 2017; Ortega et al. 2019). The confidence interval for the lentil *qDTF6a* locus detected in this study also spans the orthologous *FT* cluster (**Figure 5A**) and as in the other species, there is strong evidence that the allelic difference influences expression of one or more of these genes. In two independent NIL pairs differing at the *DTF6a* locus, *FTa1* showed strong differential expression, with expression undetectable in the late-flowering *DTF6a* genotypes but evident in the early-flowering *dtf6a* (ILL 2601) genotypes. *FTa2* was expressed in *DTF6a* lines and showed only weakly differential expression, while *FTc* expression was negligible in all lines. In addition, none of the other three *FT* genes in other genomic locations were differentially expressed (**Figure 5B**).

Functional evidence from pea and Medicago implicates *FTa1* as the most important of the three genes in the cluster. In both species, deleterious polymorphisms within the coding region of *FTa1* result in late flowering, while overexpression in Medicago confers early flowering and reduced sensitivity to both photoperiod and vernalization (Hecht et al. 2011; Laurie et al. 2011). Although we cannot definitively exclude a contribution of *FTa2* to the observed effect of the lentil *DTF6a* locus, the expression change was much weaker and seen only in one NIL pair. In addition, relative to *FTa1* genes, the pea and Medicago *FTa2* genes have minimal capacity to complement an *Arabidopsis ft* mutant (Laurie et al. 2011; Hecht et al. 2011), and in chickpea, lines with a complete deletion of *FTa2* did not display a significant difference in flowering behaviour (Ortega et al. 2019).

The fact that the early *DTF6a* allele from ILL 2601 shows dominant inheritance and confers increased *FTa1* gene expression is consistent with a scenario in which the normal repression of *FTa1* under non-inductive (i.e. short-day and/or unvernalized) conditions has been compromised. In other species, early-flowering variants co-locating with *FT* genes have been variously attributed to alterations in the promoter or other regulatory regions, tandem duplication or increase in copy number, or gain-of-function missense mutations (e.g., Beales et al. 2007; Nitcher et al. 2013). In narrow-leafed lupin (*Lupinus angustifolius*), dominant alleles at the *Ku* locus confer early, vernalization insensitive-flowering and increased expression of the single underlying *FTc*-clade gene, and feature a deletion within its promoter that presumably harbours important repressive elements (Nelson et al. 2017; Taylor et al. 2018). In addition, in Medicago, retroelement insertions in the third intron or in the 3’ UTR of *FTa1* confer a similar dominantly-inherited early-flowering phenotype, and allow expression of *FTa1* to occur in the absence of vernalization (Jaudal et al. 2013). While this could possibly reflect a direct activation by the insertion it might also represent interference with a repressive mechanism. Among many sequence differences across the *FTa1-FTa2* region (**Supp Figure 7**), the most prominent was a 7.4 kb deletion in the *FTa1*-*FTa2* intergenic region (**Figure 6**) which might plausibly harbour repressive elements acting on *FTa1*. The incidence of this polymorphism in a diversity panel capturing most of the genetic and geographic diversity within cultivated lentil germplasm (Wright et al. 2021) suggests that the *dtf6a* allele is largely restricted to, and strongly enriched in South Asian germplasm. Although this might partly be attributable to a founder effect, it also seems likely to reflect positive selection, and implies that *DTF6a* may contribute to earliness in the *pilosae* ecotype more generally.

In a recent study parallel to this one, interspecific genetic analysis of flowering time differences between *L. culinaris* and its putative wild ancestor *L. orientalis* also revealed a QTL co-locating with *qDTF6a*, and identified *FTa1* as the only differentially regulated candidate gene within the QTL interval (Yuan et al. 2021). Interestingly in this case, *FTa1* expression was higher, and flowering was earlier, in the wild parent relative to the domesticated parent cv. Lupa. This implies the existence of another derived variant distinct from that in ILL 2601, potentially involving loss-rather than gain-of-function. Together these results suggest that regulatory and structural changes that alter *FTa1* expression are an important component of natural variation for flowering time in lentil.

### Genetic control of flowering time component phases

The period between sowing and emergence, known as the pre-emergent phase (Roberts et al. 1986) has typically been subsumed within flowering time measurements based on sowing date. However, our genetic analysis identified two loci that influence this interval, that are located in genomic regions independent of those controlling the time from emergence to flowering. Broadly speaking, the pre-emergent phase itself also has several physiological components, incorporating physical and physiological dormancy and early shoot growth. This study excludes any contribution from physical dormancy since all seed coats were scarified. Interestingly, the tight co-location of a QTL for DTE with those for internode length and plant height on chromosome 7 suggests that these traits might all reflect the action of a gene involved in growth rate of the main stem, which would lead to a faster emergence of the shoot, longer internodes and thus a taller plant.

We also observed differences in the period between flower initiation and flower opening. In general, under LD conditions, the first flower to initiate is also the first to develop and open, but under SD, flower buds once formed may then fail to develop for a number of nodes (**Figure 1E**). This phenomenon is observed in a number of other temperate legumes, including pea and chickpea (Murfet, 1985; Roberts et al. 1985), and exposes a disjunction between the genetic program specifying the developmental transition to flowering, and the more general physiological orientation of the plant towards reproduction. Interestingly, we were able to associate this tendency specifically to the *DTF6a* region (**Figure 3**). Thus, while both *DTF6a* and *DTF6b* govern the node and time of first open flower under SD, the ILL 2601 allele at *DTF6a* also has an additional effect promoting the early formation of flower buds. These observations are potentially significant because the arrest of floral buds may be one consequence of exposure to environmental stresses in pulse crops, such as cool temperatures in chickpea (Nayyar et al. 2005a; Nayyar et al. 2005b; Fang et al. 2010). An improved knowledge of its genetic control and its relationship to phenology more generally may help in understanding and managing it.

## REFERENCES

Beales J, Turner A, Griffiths S, Snape J, Laurie D (2007) A Pseudo-Response Regulator is misexpressed in the photoperiod insensitive Ppd-D1a mutant of wheat (Triticum aestivum L.). Theoretical and Applied Genetics 115 (5):721–733. doi:10.1007/s00122-007-0603-4

Churchill GA, Doerge RW (1994) Empirical threshold values for quantitative trait mapping. Genetics 138:963–971

Erskine W, Adham Y, Holly L (1989) Geographic distribution of variation in quantitative traits in a world lentil collection. Euphytica 43 (1):97–103. doi:10.1007/BF00037901

Erskine W, Chandra S, Chaudhry M, Malik IA, Sarker A, Sharma B (1998) A bottleneck in lentil: widening its genetic base in South Asia. Euphytica 101:207–211

Erskine W, Ellis R, Summerfield R, Roberts E, Hussain A (1990a) Characterization of responses to temperature and photoperiod for time to flowering in a world lentil collection. Theoretical and Applied Genetics 80:193–199

Erskine W, Hussain A, Tahir M, Bahksh A, Ellis R, Summerfield R (1994) Field evaluation of a model of photothermal flowering responses in a world lentil collection. Theoretical and Applied Genetics 88:423–428

Erskine W, Saxena MC (1993) Lentil in South Asia. In: Proceedings of the Seminar on Lentil in South Asia, 11-15th March 1991, New Delhi, India. ICARDA

Fang X, Turner NC, Yan G, Li F, Siddique KH (2010) Flower numbers, pod production, pollen viability, and pistil function are reduced and flower and pod abortion increased in chickpea (Cicer arietinum L.) under terminal drought. J Exp Bot 61 (2):335–345

Food and Agricultural Organization of the United Nations (2020) FAOSTAT. http://www.fao.org/faostat/en/#data/QC. Accessed 15th April 2020

Gujaria-Verma N, Vail SL, Carrasquilla-Garcia N, Penmetsa RV, Cook DR, Farmer AD, Vandenberg A, Bett KE (2014) Genetic mapping of legume orthologs reveals high conservation of synteny between lentil species and the sequenced genomes of Medicago and chickpea. Front Plant Sci 5:676. doi:10.3389/fpls.2014.00676

Hecht V, Laurie RE, Vander Schoor JK, Ridge S, Knowles CL, Liew LC, Sussmilch FC, Murfet IC, Macknight RC, Weller JL (2011) The pea GIGAS gene is a FLOWERING LOCUS T homolog necessary for graft-transmissible specification of flowering but not for responsiveness to photoperiod. Plant Cell 23 (1):147–161. doi:10.1105/tpc.110.081042

Hodges T (1990) Predicting crop phenology. CRC Press, Boca Raton, Florida

Iqbal A, Khalil IA, Ateeq N, Khan MS (2006) Nutritional quality of important food legumes. Food Chemistry 97:331–335

Jaudal M, Yeoh CC, Zhang LL, Stockum C, Mysore KS, Ratet P, Putterill J (2013) Retroelement insertions at the Medicago FTa1 locus in spring mutants eliminate vernalisation but not long-day requirements for early flowering. Plant J 76 (4):580–591. doi:Doi 10.1111/Tpj.12315

Keatinge J, Qi A, Kusmenoglu I, Ellis R, Summerfield R, Erskine W (1995) Defining critical weather events in the phenology of lentil for winter sowing in the west Asian highlands. Agricultural and forest meteorology 74:251–263

Keatinge J, Qi A, Kusmenoglu I, Ellis R, Summerfield R, Erskine W (1996) Using genotypic variation in flowering responses to temperature and photoperiod to select lentil for the west Asian highlands. Agricultural and forest meteorology 78:53–65

Kumar J, Gupta S, Gupta P, Dubey S, Tomar R, Kumar S (2016) Breeding strategies to improve lentil for diverse agro-ecological environments. Indian Journal of Genetics and Plant Breeding 76:530–549

Laurie RE, Diwadkar P, Jaudal M, Zhang LL, Hecht V, Wen JQ, Tadege M, Mysore KS, Putterill J, Weller JL, Macknight RC (2011) The Medicago FLOWERING LOCUS T Homolog, MtFTa1, Is a Key Regulator of Flowering Time. Plant Physiology 156 (4):2207–2224. doi:DOI 10.1104/pp.111.180182

Lin X, Liu B, Weller JL, Abe J, Kong F (2021) Molecular mechanisms for the photoperiodic regulation of flowering in soybean. J Integr Plant Biol 63 (6):981–994

Maqbool A, Shafiq S, Lake L (2010) Radiant frost tolerance in pulse crops-a review. Euphytica 172 (1):1–12. doi:10.1007/s10681-009-0031-4

Murfet IC (1985) Pisum sativum. In: Halevy AH (ed) CRC handbook of flowering, vol IV. CRC Press, pp 97–126

Nayyar H, Bains T, Kumar S (2005a) Low temperature induced floral abortion in chickpea: Relationship to abscisic acid and cryoprotectants in reproductive organs. Environmental and Experimental Botany 53 (1):39–47. doi:10.1016/j.envexpbot.2004.02.011

Nayyar H, Bains TS, Kumar S, Kaur G (2005b) Chilling effects during seed filling on accumulation of seed reserves and yield of chickpea. J Sci Food Agric 85 (11):1925–1930. doi:10.1002/jsfa.2198

Nelson MN, Ksiazkiewicz M, Rychel S, Besharat N, Taylor CM, Wyrwa K, Jost R, Erskine W, Cowling WA, Berger JD, Batley J, Weller JL, Naganowska B, Wolko B (2017) The loss of vernalization requirement in narrow-leafed lupin is associated with a deletion in the promoter and de-repressed expression of a Flowering Locus T (FT) homologue. New Phytol 213 (1):220–232. doi:10.1111/nph.14094

Nitcher R, Distelfeld A, Tan C, Yan L, Dubcovsky J (2013) Increased copy number at the HvFT1 locus is associated with accelerated flowering time in barley. Mol Genet Genomics 288 (5-6):261–275. doi:10.1007/s00438-013-0746-8

Ortega R, Hecht VFG, Freeman JS, Rubio J, Carrasquilla-Garcia N, Mir RR, Penmetsa RV, Cook DR, Millan T, Weller JL (2019) Altered Expression of an FT Cluster Underlies a Major Locus Controlling Domestication-Related Changes to Chickpea Phenology and Growth Habit. Front Plant Sci 10 (824). doi:10.3389/fpls.2019.00824

Ramsay L, Koh CS, Kagale S, Gao D, Kaur S, Haile T, Gela TS, Chen L-A, Cao Z, Konkin DJ, Toegelová H, Doležel J, Rosen BD, Stonehouse R, Humann JL, Main D, Coyne CJ, McGee RJ, Cook DR, Varma Penmetsa R, Vandenberg A, Chan C, Banniza S, Edwards D, Bayer PE, Batley J, Udupa SM, Bett KE (2021) Genomic rearrangements have consequences for introgression breeding as revealed by genome assemblies of wild and cultivated lentil species. bioRxiv:2021.2007.2023.453237. doi:10.1101/2021.07.23.453237

Roberts E, Hadley P, Summerfield R (1985) Effects of temperature and photoperiod on flowering in chickpeas (Cicer arietinum L.). Annals of Botany 55:881–892

Roberts EH, Summerfield RJ, Muehlbauer FJ, Short RW (1986) Flowering in Lentil (Lens culinaris Medic.): The Duration of the Photoperiodic Inductive Phase as a Function of Accumulated Daylength above the Critical Photoperiod. Annals of Botany 58 (2):235–248. doi:10.1093/oxfordjournals.aob.087201

Sansaloni C, Petroli C, Jaccoud D, Carling J, Detering F, Grattapaglia D (2011) Diversity Arrays Technology (DArT) and next-generation sequencing combined: genome-wide, high throughput, highly informative genotyping for molecular breeding of Eucalyptus. BMC Proceedings 5:1–2

Sarker A, Erskine W, Sharma B, Tyagi M (1999) Inheritance and linkage relationship of days to flower and morphological loci in lentil (Lens culinaris Medikus subsp. culinaris). Journal of Heredity 90:270–275

Sharpe AG, Ramsay L, Sanderson L-A, Fedoruk MJ, Clarke WE, Li R, Kagale S, Vijayan P, Vandenberg A, Bett KE (2013) Ancient orphan crop joins modern era: gene-based SNP discovery and mapping in lentil. BMC Genomics 14 (1):192. doi:10.1186/1471-2164-14-192

Sita K, Sehgal A, HanumanthaRao B, Nair RM, Vara Prasad PV, Kumar S, Gaur PM, Farooq M, Siddique KHM, Varshney RK, Nayyar H (2017) Food Legumes and Rising Temperatures: Effects, Adaptive Functional Mechanisms Specific to Reproductive Growth Stage and Strategies to Improve Heat Tolerance. Front Plant Sci 8 (1658). doi:10.3389/fpls.2017.01658

Summerfield R, Roberts E, Erskine W, Ellis R (1985) Effects of temperature and photoperiod on flowering in lentils (Lens culinaris Medic.). Annals of Botany 56:659–671

Sussmilch FC, Berbel A, Hecht V, Vander Schoor JK, Ferrándiz C, Madueño F, Weller JL (2015) Pea VEGETATIVE2 is an FD homolog that is essential for flowering and compound inflorescence development. Plant Cell 27 (4):1046–1060. doi:10.1105/tpc.115.136150

Tang H, Krishnakumar V, Bidwell S, Rosen B, Chan A, Zhou S, Gentzbittel L, Childs KL, Yandell M, Gundlach H, Mayer KF, Schwartz DC, Town CD (2014) An improved genome release (version Mt4.0) for the model legume Medicago truncatula. BMC Genomics 15 (1):312. doi:10.1186/1471-2164-15-312

Taylor CM, Kamphuis LG, Zhang W, Garg G, Berger JD, Mousavi-Derazmahalleh M, Bayer PE, Edwards D, Singh KB, Cowling WA, Nelson MN (2018) INDEL variation in the regulatory region of the major flowering time gene LanFTc1 is associated with vernalization response and flowering time in narrow-leafed lupin (Lupinus angustifolius L.). Plant Cell Environ

Van Ooijen J (2006) JoinMap® 4, Software for the calculation of genetic linkage maps in experimental populations. Kyazma BV, Wageningen, The Netherlands

Van Ooijen JW (2009) MapQTL 6: Software for the mapping of quantitative trait loci in experimental populations of diploid species (Kyazma B.V., Wageningen)

Voorrips R (2002) MapChart: software for the graphical presentation of linkage maps and QTLs. Journal of Heredity 93:77–78

Weller JL, Liew LC, Hecht VF, Rajandran V, Laurie RE, Ridge S (2012) A conserved molecular basis for photoperiod adaptation in two temperate legumes. Proceedings of the National Academy of Sciences 109:21158–21163

Weller JL, Ortega R (2015) Genetic control of flowering time in legumes. Frontiers in Plant Science 6

Weller JL, Vander Schoor JK, Perez-Wright EC, Hecht V, González AM, Capel C, Yuste-Lisbona FJ, Lozano R, Santalla M (2019) Parallel origins of photoperiod adaptation following dual domestications of common bean. Journal of Experimental Botany 70 (4):1209–1219

Wickland, D. P., and Hanzawa, Y. (2015). The FLOWERING LOCUS T / TERMINAL FLOWER 1 gene family: functional evolution and molecular Mechanisms. Mol. Plant 8, 983–997

Wright DM, Neupane S, Heidecker T, Haile TA, Chan C, Coyne CJ, McGee RJ, Udupa S, Henkrar F, Barilli E, Rubiales D, Gioia T, Logozzo G, Marzario S, Mehra R, Sarker A, Dhakal R, Anwar B, Sarkar D, Vandenberg A, Bett KE (2021) Understanding photothermal interactions will help expand production range and increase genetic diversity of lentil (Lens culinaris Medik.). PLANTS, PEOPLE, PLANET 3 (2):171–181. doi:https://doi.org/10.1002/ppp3.10158

Yuan HY, Caron CT, Ramsay L, Fratini R, Pérez de la Vega M, Vandenberg A, Weller JL, Bett KE (2021) Genetic and gene expression analysis of flowering time regulation by light quality in lentil. Ann Bot. doi:10.1093/aob/mcab083

