## Supplementary Material for "Genetic analysis of early phenology in lentil identifies distinct loci controlling component traits"

**Supplementary Table 1.** List of all accessions used in Figure 1.

| Accession | Type[a] | Other name(s) | Origin | Latitude[b] | Comments |
|---|---|---|---|---|---|
| ILWL 7 | Wild | - | Turkey | 38°N | |
| ILL 119 | LR | PI 170478 | Turkey | 37°N | |
| ILL 131 | LR | PI 174251 | Turkey | 38°N | |
| ILL 798 | LR | F75 | Egypt | 30°N | |
| ILL 1702 | LR | EL32 | Ethiopia | 11°N | |
| ILL 1744 | LR | EL 79 | Ethiopia | 10°N | |
| ILL 1756 | LR | 72L | Afghanistan | 35°N | |
| ILL 1771 | LR | 326L | Afghanistan | 35°N | |
| ILL 2601 | LR | LWS 16 | India | 23°N | |
| ILL 4349 | CV | Laird | Russia | 50°N | |
| ILL4400 | LR | Syrian local large | Syra | 36°N | |
| ILL 4605 | CV | Precoz | Argentina | 27°S | |
| ILL 5584 | | 78526004 | Jordan | 32°N | Selection from LR |
| ILL 5588 | CV | Talya 2 | Jordan | 32°N | Released in Lebanon as Talya 2 |
| ILL 6005 | | | | | Selection from cross of Laird x Precoz |
| ILL 6007 | | | | | Selection from cross of Laird x Precoz |

[a] LR = Landrace; CV = Cultivar; Wild = *Lens orientalis*
[b] approximate latitude of origin

**Supplementary Table 2.** List of primers used in this study for gene expression analysis.

| Target gene | Primer name | Sequence | Reference |
|---|---|---|---|
| *FTa1* | LcFTa1-8F | CCGATATTCCAGCAACTACTGA | |
| | LcFTa1-7R | AACACGAACACGAAACGATG | |
| *FTa2* | LcFTa2-F2 | ACCCCATGATCTACTACACCC | |
| | LcFTa2-4R | CCATCACAATTCAAAGCAATG | |
| *FTa3* | LcFTa3-F1 | AGTTCCAGGAATATCAGTCACC | |
| | LcFTa3-R1 | CCAAGGGTTAAGGTTGGTGG | |
| *FTb2* | LcFTb-3F | GGTGAACCCTGATGCACCTA | |
| | LcFTb-1R | GAACGTTGTCCCAGTAGTCG | |
| *FTc* | LcFTc-F3 | GGTGCATCTGCGTCCACC | |
| | LcFTc-R3 | ACAATTGGTTAATCGTCCAAGGG | |
| *Actin* | LcActin-Fw | CCAAATCATGTTTGAGGCTTTTAA | Sen Gupta et al., 2017 |
| | LcActin-Rv | GTGAAAGAACGGCCTGAATAGC | |

**Supplementary Table 3.** Sequence associated with DArT markers in the ILL 2601 x ILL 5588 $F_2$ genetic linkage map.

Supplementary Table 3 is offered in an independent excel file.

**Supplementary Table 5** Mean (µ) and standard deviation values obtained for days from emergency to first flower (DTF) and node of the first open flower (NFD) in lentil lines ILL 2601 and ILL 5588 grown in 4 different conditions.

| | | | DTF | | NFD | |
|---|---|---|---|---|---|---|
| **Accession** | **Treatment** | **N** | **Mean** | **Std. Dev** | **Mean** | **Std. Dev** |
| **ILL 2601** | **NV-LD** | 12 | 35.5 | 2.5 | 8.2 | 0.9 |
| | **V-LD** | 19 | 37.9 | 3.3 | 8.0 | 1.0 |
| | **NV-SD** | 9 | 65.7 | 5.7 | 10.8 | 0.8 |
| | **V-SD** | 10 | 62.8 | 8.6 | 10.6 | 1.3 |
| **ILL 5588** | **NV-LD** | 13 | 41.4 | 2.8 | 13.6 | 1.2 |
| | **V-LD** | 21 | 39.1 | 2.5 | 12 | 1.5 |
| | **NV-SD** | 15 | -[a] | -[a] | -[a] | -[a] |
| | **V-SD** | 23 | 71.8 | 4.8 | 13 | 1.4 |

N= Number of plants; LD = Long days; SD = Short days; V = Vernalized plants; NV= Non-vernalized plants

-[a] Plants were unable to flower/set pods under this conditions during the scoring period.

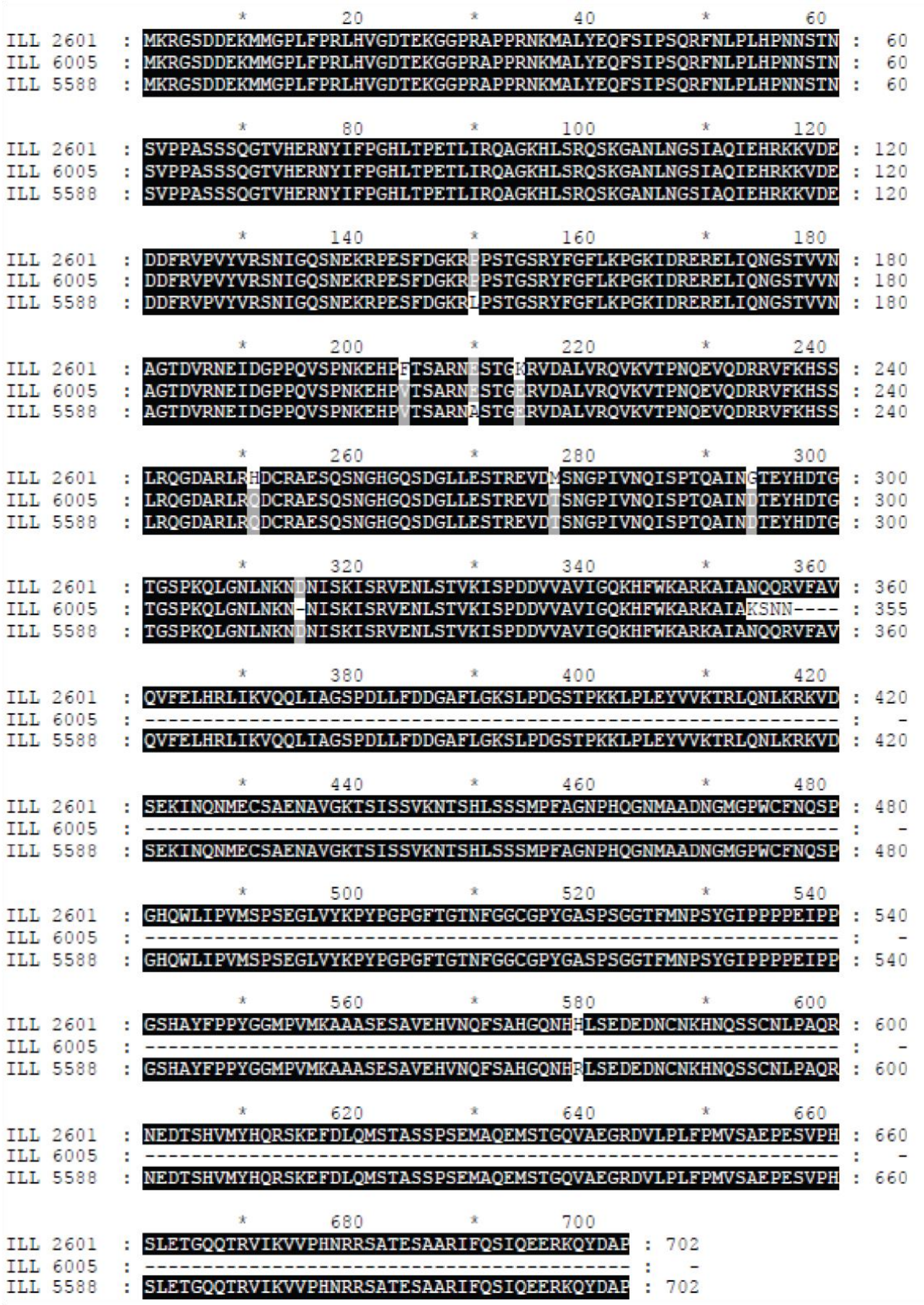


**Supplementary Figure 1.** Multiple sequence alignment of the ELF3 proteins in lentil accessions ILL 6005, ILL 5588 and ILL 2601. Shades indicate minimum conservation level: Black = 80%, grey = 60%.

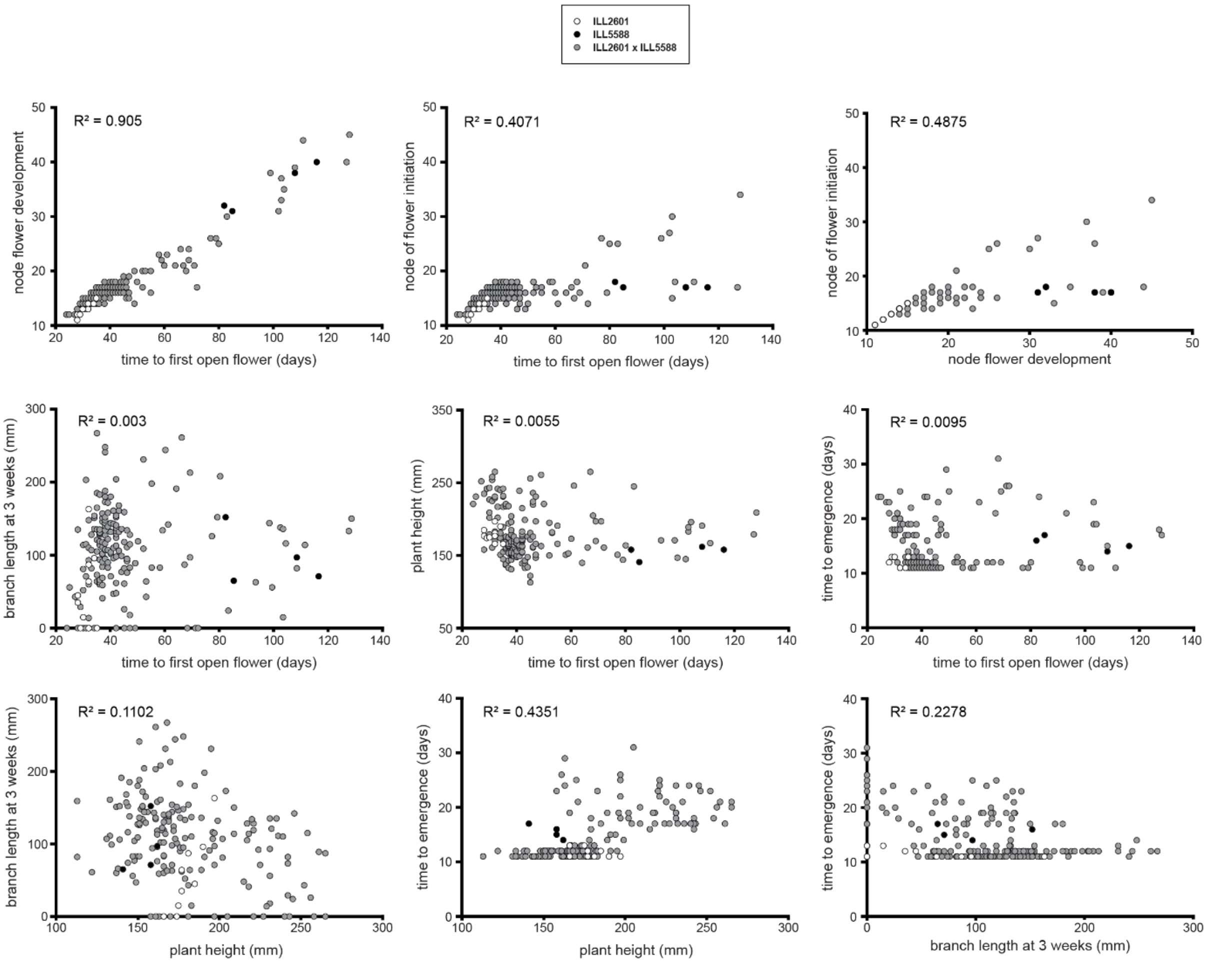


**Supplementary Figure 2.** Correlation of traits measured in the ILL 2601 x ILL 5588 $F_2$ mapping population. The reduced correlation of floral initiation with flower development and flowering time is indicative of flower abortion in some lines.

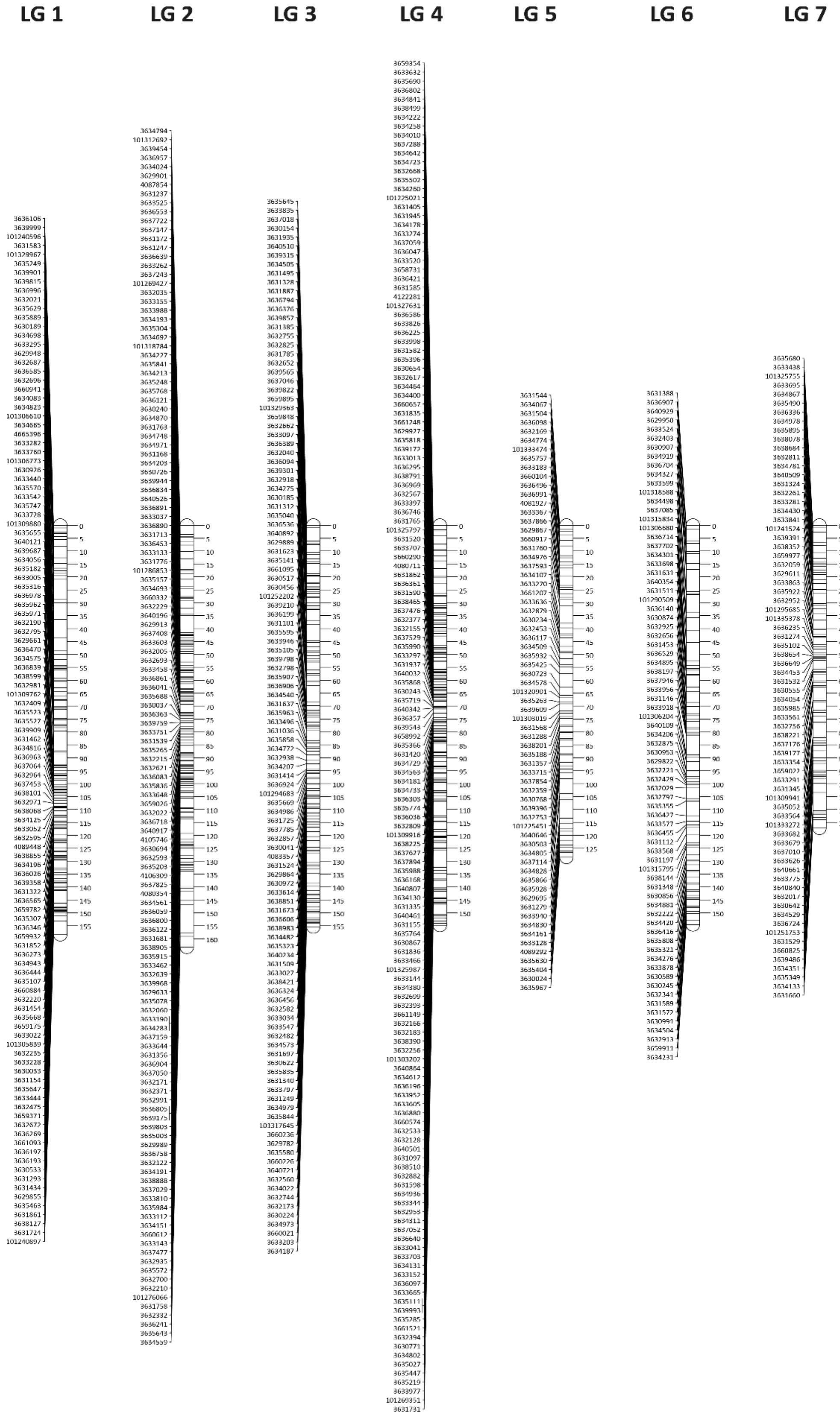


**Supplementary figure 3. ILL 2601 x ILL 5588 F2 genetic linkage map**

Genetic linkage map consists of seven linkage groups corresponding to the seven chromosomes of the *Lens* genus. The nomenclature proposed is adapted from Sharpe et al. (2013).

**Supplementary Table 4.** Statistics of the genetic linkage map displayed in Supplementary Figure 3.

| Linkage group | Markers | Length (cM) | Density (markers/cM) |
|---|---|---|---|
| 1 | 115 | 158.40 | 1.38 |
| 2 | 136 | 163.35 | 1.20 |
| 3 | 118 | 155.43 | 1.32 |
| 4 | 151 | 154.81 | 1.03 |
| 5 | 67 | 128.51 | 1.92 |
| 6 | 75 | 154.63 | 2.06 |
| 7 | 72 | 117.16 | 1.63 |
| TOTAL | 734 | 1032.28 | 1.41 |

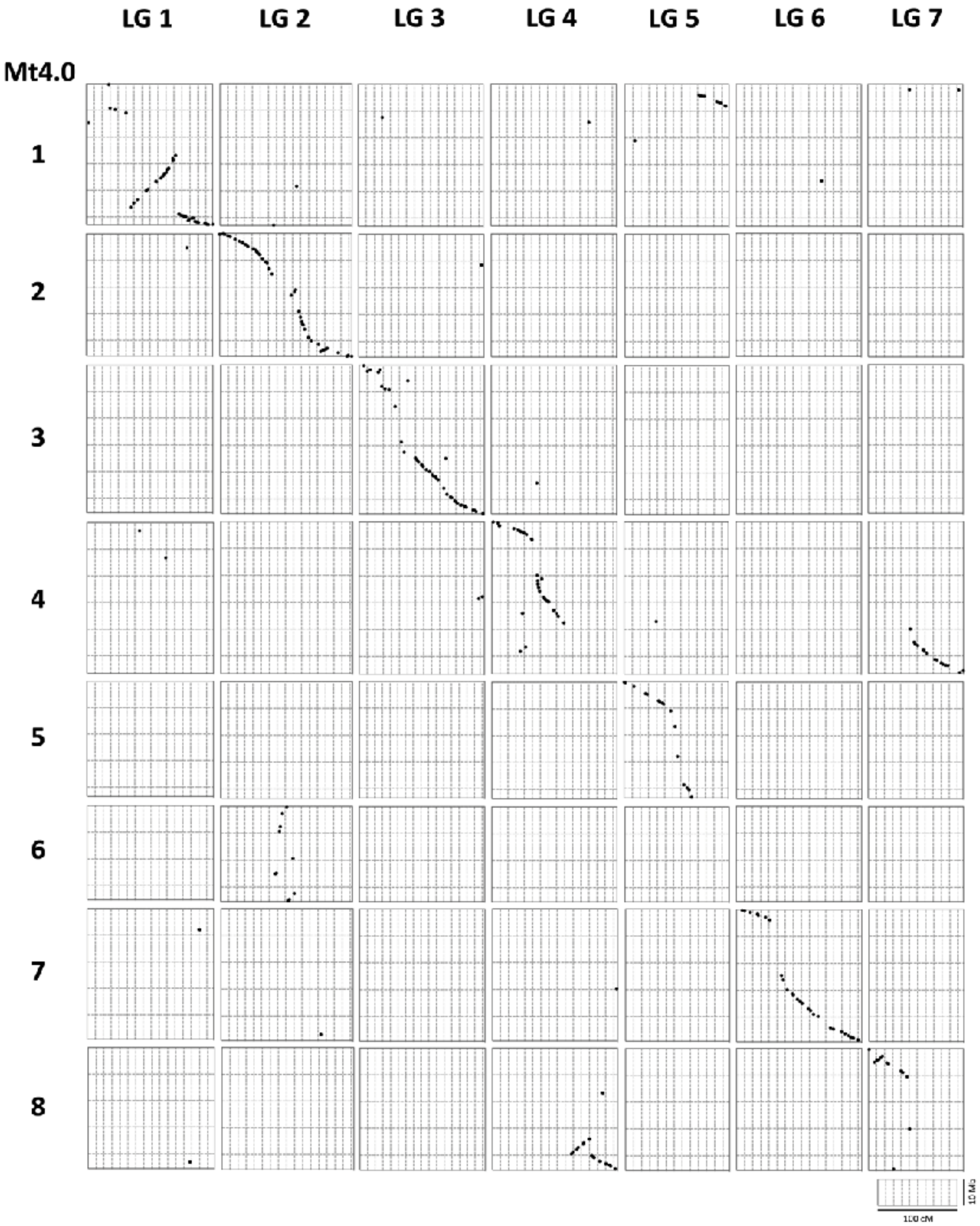


**Supplementary figure 4. Synteny between lentil genetic linkage map and *M. truncatula* genome (Mt4.0)**

Each Y-axis interval represents 10 Mb in the Mt4.0 assembly and each X-axis interval represents 10 cM in the linkage map. 256 markers with sequences demonstrating significant similarities (*e*-value < 0.001), and 33 markers (*e*-value < 0.1) and 38 markers (*e*-value < 0.01) with sequences demonstrating low similarities are visualised in the dot plot. The dot plot was manually graphed using Prism 6.

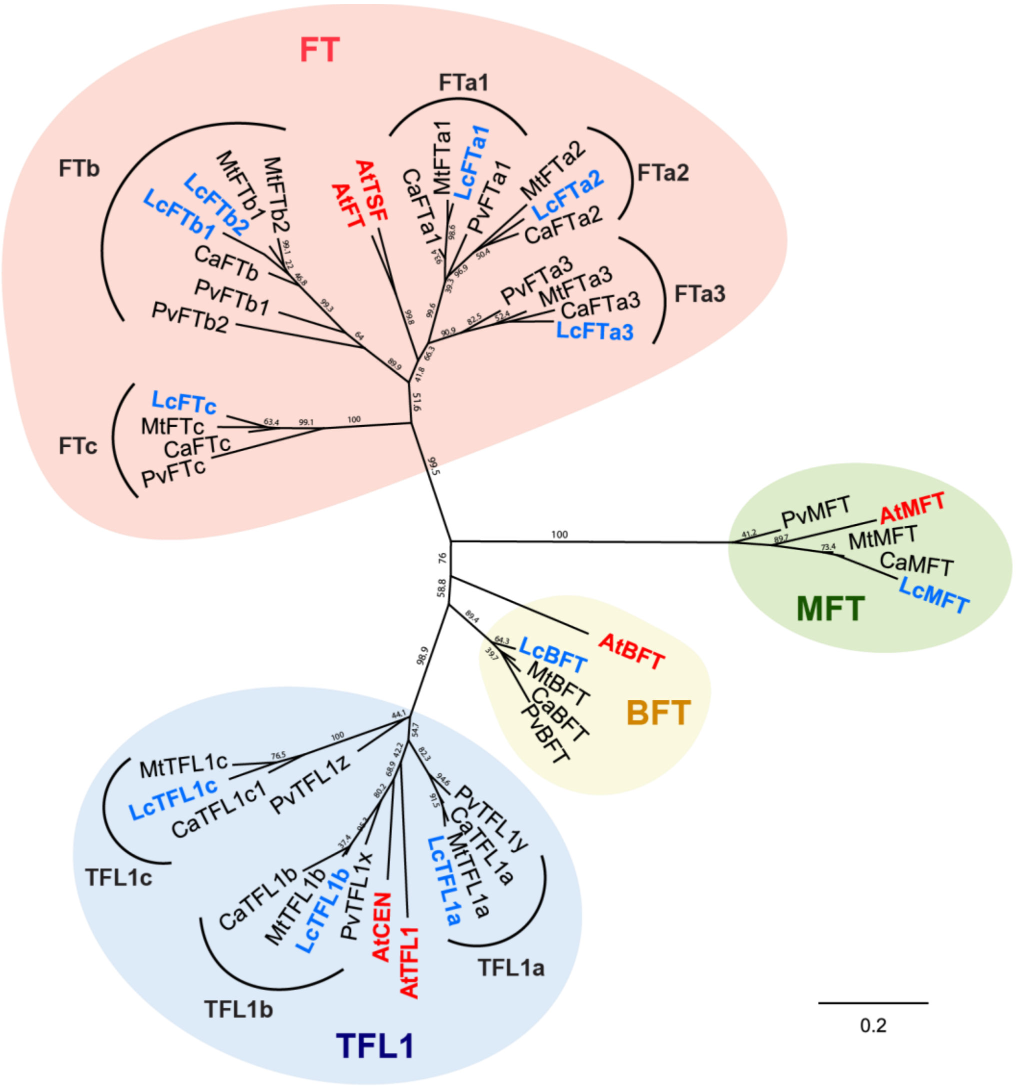


**Supplementary Figure 5.** Phylogenetic tree of the phosphatidyl ethanolamine-binding protein (PEBP) gene family in five legume species (Lc=*Lens culinaris*, in blue, Mt = *Medicago truncatula*, Ca = *Cicer arietinum*, Pv = *Phaseolus vulgaris*) and *Arabidopsis thaliana* (At, in red). Protein sequences from gene accessions in supplementary table 6 were aligned in Geneious Prime 2020.2.2 (http://www.geneious.com) using MAFFT. Phylogenetic tree was inferred using PhyML .3.20180621 (Guindon et al. 2010) , Blosum62 method and a bootstraping of 1000 replications.

**Supplementary table 6.** Gene accession numbers of the 48 PEBP amino acid sequences from five plant species used to build the phylogenetic tree displayed in supplementary figure 6.

| **Arabidopsis** | |
|---|---|
| **Gene** | **Accession** |
| ***Arabidopsis thaliana*** | |
| AtBFT | AT5G62040 |
| AtCEN | AT2G27550 |
| AtFT | AT1G65480 |
| AtMFT | AT1G18100 |
| AtTFL1 | AT5G03840 |
| AtTSF | AT4G20370 |

| **Legumes** | | | |
|---|---|---|---|
| **Gene** | **Accession** | **Gene** | **Accession** |
| ***Phaseolus vulgaris*** | | ***Cicer arietinum*** | |
| PvBFT | Phvul.004G119700 | CaBFT | LOC101507903 |
| PvFTa1 | Phvul.001G097300 | CaFTa1 | LOC101497376 |
| PvFTa3 | Phvul.004G074700 | CaFTa2 | LOC101496618 |
| PvFTb1 | Phvul.008G003800 | CaFTa3 | LOC101515383 |
| PvFTb2 | Phvul.008G003700 | CaFTb | LOC101505276 |
| PvFTc | Phvul.001G097200 | CaFTc | LOC101508200 |
| PvMFT | Phvul.002G327700 | CaMFT | LOC101504081 |
| PvTFL1x | Phvul.005G124600 | CaTFL1a | LOC101506075 |
| PvTFL1y | Phvul.001G189200 | CaTFL1b | LOC101508699 |
| PvTFL1z | Phvul.007G229300 | CaTFL1c1 | LOC101495644 |
| ***Lens culinaris*** | | ***Medicago truncatula*** | |
| LcBFT | Lcu.2RBY.2g038980 | MtBFT | Medtr0020s0120 |
| LcFTa1 | Lcu.2RBY.6g043850 | MtFTa1 | Medtr7g084970 |
| LcFTa2 | Lcu.2RBY.6g043870 | MtFTa2 | Medtr7g085020 |
| LcFTa3 | Lcu.2RBY.2g046590 | MtFTa3 | Medtr6g033040 |
| LcFTb1 | Lcu.2RBY.6g000730 | MtFTb1 | Medtr7g006630 |
| LcFTb2 | Lcu.2RBY.6g000760 | MtFTb2 | Medtr7g006690 |
| LcFTc | Lcu.2RBY.6g043940 | MtFTc | Medtr7g085040 |
| LcMFT | Lcu.2RBY.4g081710 | MtMFT | Medtr8g106840 |
| LcTFL1a | Lcu.2RBY.6g060590 | MtTFL1a | Medtr7g104460 |
| LcTFL1b | Lcu.2RBY.2g084140 | MtTFL1b | Medtr2g086270 |
| LcTFL1c | Lcu.2RBY.1g033180 | MtTFL1c | Medtr1g060190 |

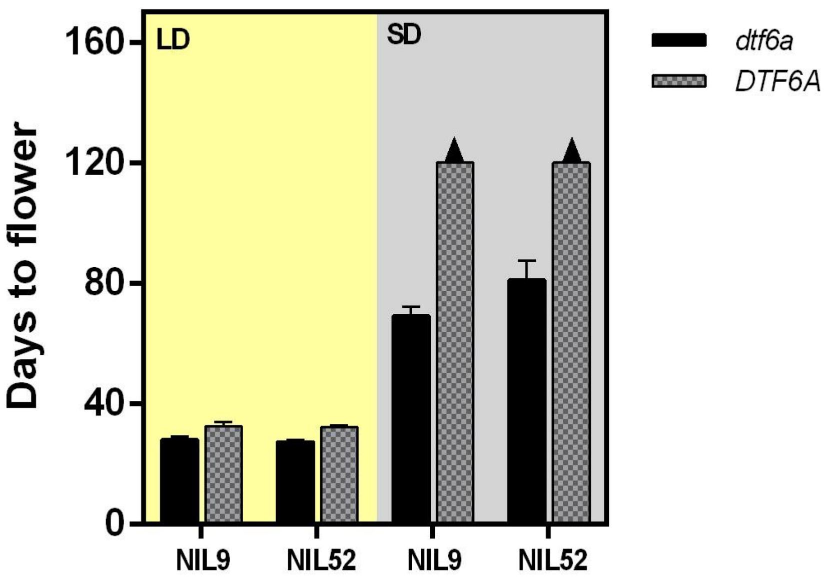


**Supplementary Figure 6.** Flowering phenotype of two pair of NILs segregating for *DTF6a*, evaluated under LD (16 hours photoeriod, yellow background) and SD (8 hours photoperiod, grey background) conditions. Arrows over the columns indicate that plants were unable to flower after 120 days.

**Supplementary Figure 7.** Graphical overview of the alignment between the *FTa1-FTa2* sequences from lentil accessions ILL 5588 and ILL 2601. Grey boxes over the alignment indicate the extent of *FTa1* and *FTa2* genes. Yellow triangles represent exons and yellow lines introns. Sequence similarities are shown in grey and polymorphisms in black. Relevant deletions are highlighted in red.

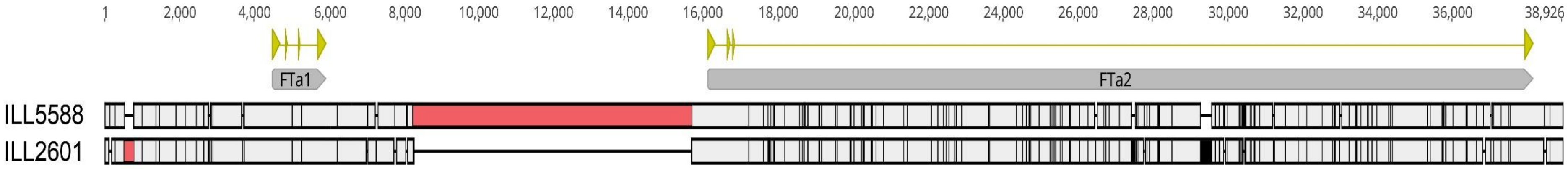


**Supplementary table 7.** Summary of all the polymorphisms found in the comparison between the *FTa1-FTa2* sequences from lentil accessions ILL 5588 and ILL 2601.

| | FTa1 5' UTR | FTa1 Gene | Intergenic | FTa2 Gene | FTa2 3' UTR | Total |
|---|---|---|---|---|---|---|
| Position (alignment) | 1-4491 | 4492-5925 | 5926-16117 | 16118-38173 | 38174-38926 | |
| Size (alignment) | 4491 | 1434 | 10192 | 22056 | 753 | 38926 |
| Size (ILL 5588) | 4230 | 1434 | 10173 | 21652 | 753 | 38242 |
| Size (ILL 2601) | 4489 | 1434 | 2708 | 21974 | 747 | 31352 |
| SNPs | 14 | 3 | 5 | 113 | 1 | 136 |
| Indels | 5 | 0 | 5 | 14 | 1 | 25 |
| Total | 19 | 3 | 10 | 127 | 2 | 161 |

**Supplementary table 8.** Description of all the SNPs found in the 38926 bp-length alignment of the *FTa1-FTa2* sequences from lentil accessions ILL 5588 and ILL 2601. For each SNP, the first variant given in the table correspond to the allele present in ILL 5588 and the second to that from ILL 2601.

| Position | SNP | Position | SNP | Position | SNP | Position | SNP | Position | SNP | Position | SNP | Position | SNP | Position | SNP |
|---|---|---|---|---|---|---|---|---|---|---|---|---|---|---|---|
| 295 | G/T | 7040 | A/T | 19919 | T/C | 22681 | C/T | 25765 | A/T | 28500 | T/G | 32822 | C/T | 35816 | T/C |
| 782 | A/G | 8084 | A/G | 19938 | C/T | 22734 | C/T | 25781 | T/C | 29655 | T/C | 32846 | C/T | 36016 | A/G |
| 990 | C/A | 8085 | T/A | 20002 | C/A | 22878 | C/T | 26049 | A/G | 29982 | T/A | 33235 | A/T | 36347 | A/G |
| 1362 | G/A | 8086 | T/A | 20191 | C/T | 23592 | T/C | 26258 | A/C | 30294 | G/A | 33405 | T/C | 36374 | G/C |
| 1454 | G/T | 17191 | C/A | 20249 | T/A | 23633 | C/T | 26682 | C/T | 30331 | C/T | 33434 | C/T | 36637 | T/C |
| 1904 | G/T | 17605 | C/T | 20254 | T/C | 24343 | C/T | 26719 | T/C | 30362 | C/T | 33449 | A/G | 37102 | A/G |
| 2163 | A/G | 17801 | A/T | 20265 | C/T | 24519 | A/T | 26835 | A/T | 30466 | C/T | 33545 | C/T | 37280 | G/A |
| 2444 | G/A | 17874 | C/T | 20467 | G/C | 24599 | A/G | 26836 | G/T | 30598 | T/C | 33730 | C/T | 37493 | C/T |
| 2643 | A/G | 17897 | C/T | 20489 | C/T | 24690 | A/C | 26837 | A/G | 30619 | T/C | 33747 | A/G | 37531 | C/T |
| 2774 | A/G | 18161 | A/G | 20594 | C/A | 24726 | C/T | 27103 | A/C | 30689 | C/G | 33836 | A/C | 38597 | A/G |
| 2834 | A/T | 18564 | T/C | 20804 | A/G | 24955 | A/G | 27409 | C/G | 30723 | T/C | 33968 | T/C | | |
| 2882 | T/A | 18625 | C/T | 21328 | A/T | 25245 | C/A | 27573 | A/T | 30827 | C/T | 34279 | T/A | | |
| 2902 | G/A | 18783 | A/G | 21431 | C/G | 25304 | A/T | 27623 | C/T | 31237 | T/C | 34329 | T/C | | |
| 3728 | G/T | 19068 | A/G | 22069 | G/A | 25338 | A/G | 27827 | A/G | 31533 | A/G | 34377 | C/T | | |
| 5015 | G/C | 19100 | C/T | 22240 | A/C | 25400 | T/C | 27897 | A/C | 31866 | A/G | 35310 | G/A | | |
| 5022 | A/G | 19141 | G/T | 22405 | G/C | 25517 | T/A | 28121 | T/A | 32104 | T/C | 35396 | C/T | | |
| 5258 | T/C | 19524 | A/T | 22478 | C/T | 25524 | C/T | 28126 | A/T | 32510 | C/T | 35712 | C/T | | |
| 6214 | T/C | 19554 | C/T | 22539 | T/C | 25559 | C/T | 28161 | C/A | 32785 | T/G | 35714 | A/G | | |

**Supplementary table 9.** Description of all indels found in the 38926 bp-length alignment between the *FTa1-FTa2* sequences from lentil accessions ILL 5588 and ILL 2601. For each indel, the first variant given in the table correspond to the allele present in ILL 5588 and the second to that from ILL 2601.

| Indel | Indel start | Indel end | Length | Indel | Indel start | Indel end | Length |
|---|---|---|---|---|---|---|---|
| TT/- | 141 | 142 | 2 | -/Var3 [a] | 27410 | 27515 | 106 |
| -/A | 520 | 520 | 1 | G/- | 27759 | 27759 | 1 |
| -/Var1[a] | 535 | 779 | 245 | -/Var4 [a] | 29263 | 29548 | 286 |
| -/GGGGTGGGGGGG | 2779 | 2790 | 12 | GCGTC/- | 29755 | 29759 | 5 |
| -/TTC | 3680 | 3682 | 3 | TGTTTTAAATATAAAAACTT/- | 29878 | 29897 | 20 |
| T/- | 7012 | 7012 | 1 | Var5 [a] /- | 30386 | 30437 | 52 |
| -/T | 7262 | 7262 | 1 | -/TCT | 31210 | 31212 | 3 |
| AATAATATAGTTATTTTTTA/- | 7734 | 7753 | 20 | -/TTC | 32986 | 32988 | 3 |
| ATAT/- | 8053 | 8056 | 4 | -/AG | 35740 | 35741 | 2 |
| Var2 [a] /- | 8243 | 15683 | 7441 | TTT/- | 36817 | 36819 | 3 |
| A/- | 17716 | 17716 | 1 | -/T | 37009 | 37009 | 1 |
| -/A | 18680 | 18680 | 1 | AACAGT/- | 38440 | 38445 | 6 |
| -/TT | 26458 | 26459 | 2 | | | | |

[a] Due to their excessive length, the sequences corresponding to these variants are shown separately in the next page(s) of this document

### Variant 1 sequence

TCTCTTCAAACATTTACAAACAAACACTATTTTCAAGGTAAAAACACGGATTTTCAAAAGGTTCCAGTGGAGTACCACATATGTGAGGGGTGCTAATACCTTCTCTTCGCATAAC
CAACCTCTAAACCCTTATTTCTTTATCTTGTGGGTTTTATTTGCTCTTTTCCCATTTTCCTTTTAGAAACAATAAAAGTAAGGTGGTGACTCTTGCTTTATGAGTTAAGTTAATTAA
TAGCTTAAACTCA

### Variant 2 sequence

TTCAAACAGCAGTTGTAACATATAGTTAACTGAAGAAGCTAGGCTTCAAAGAGATGATTAAAAATTTAAGAACTAAAGAAAAATAAATAAATAAAAAACGCTAAGAAATGATA
TACAAGTGGAATCCGCAATTAAGGTTTCTATTAAAAAAAATAGAAGAAAATCATAAATAGTAGTATTAATATTTTAAAATAATGTGATGGTTAAAAAGGAAATCTATATGCTAT
TGAATAGTCATTTTCTACTAATTATTTGATGATTGTATAAACATTACATCTTATATCTATCGAGAGTAGAGCTCTGGTCTACTTTCTCAAACCATTAATCATACTACTTGATCAATG
TAGGGCTATTCAGGACCTTTTCCAAAAAATAAATAATAAACAAACAACCGCTCGCTAGTTGCTATTTATGTAGTTTAACTAAGTGCAAGAAAGCAGGAACGTGTGAAAGACCA
CATCTCTCACGCTTGTAGTTTTTTCTTCTCTCAAAAATATTTATTATTTAAAATTATAATATATTATATGTTAGAAACAAAACCCTTTGAACTAGATAATATCTTCAACTGGAGGAG
AAAAAAGAAAATATCTCCATCTCACTGTTCTGCACACAGTGGCTACTACTGTCATCTATGAAATCACCTTTCCTTGTTCGCAAGGAAATTACAGTTTCTACACAAATATTTCATCT
CATTCTAAAATCTTACTTTTCCCCCAAAACATAAACCACAGAAACAACGTGAAGATGGAAGAATGTAAATGTGAAGTGAAAAAGTTTTTCACTTAAATCAGGAACCCTAGCAAA
CCACAGTTCACTGAAAGAGACGACAGTGGCAAATTTACCTGATACCACGGCAGTTATTGTGGATGACCTCGTCCTGTTACAGTGAAAATCTGATCATGTGGCGGTTCTGTTGA
CAAATCTGTAACTGTAGTTCTGGGAAGATTTTCACCCTAGGTTTTCCTGGATCTATAGTTCTACAACGTGTTAGATTTATGACGAGGATGAGTGACTTCTGGTTTTAATTTTAGA
TATATATTTATCTGAGATCTAGTTTTGTTGTAAGGGGTGTGCTTTTGAAATCATGAAAATTTGCAGGTAAATTTAACACTGTTTGTGATTTAATAGAAAATAACTTTAACTTTTTT
TTAAAATATTGAATGATCTTTGGATTTAATCATTTCCATTTTATTCATCTGTTTTATTTATTTATTTATTTAATTACGACGTGCTTTGCTTTATAGTTTTTTTCTTTATAGGGTTAAAT
TTGGCATTAAATTGTGGTTATCCAATTCCAGATCGATCTACTGGAAGGTATTTTACTTCCTAAAATTTCTATAGTTAAGTTTACACTCCTAATGTTGGCTTGGTGATTTAGTTTTG
TCCTATAGTTTAGTCTTTTTACACTGAATCTATAATGATTTAATTCTAATTATAACTACTTTTATAATTGTATAAATAAGAATAATTGTGATTTATTTGTAGTATAGAAATTTTTGCA
GTGCTTAACACACGCTAAAGCATGATCATATATATGTTAATTTGATTAATCAGTTAATTGTATTTCTGCTAGCTGTCAAATATTTTTATTACTACTATATCATTTTAAGTTTCAATT
TAATGGTTGTATGAATAAAATCTAATGTTCATTAATATATTATTATTATTATTATTATTATTATTATTATTATTATTATTATTATTATTATTATTATTATTATTATAAAGTTTAGAAGT
TATTTACATTTTATCCAAAAAAGAAGGAAGCTTACTTACATCAATTGAAATAAGAGTTGTCTATGCTCAAACTCAAATCATTAGTCATTCTCATTCAGGTCCAATTGTTAGGGAT
AGTCTGGTACTTTAACAATGAAAAAAAGAAATAAGTGCTGCCCGCTGCCCGCTGGTGCTTCTATATTAATATTAACCAATTGCAGAAACAAAGAAAAGCCGGAACATGTGCAA
TACCATATCTTCATCCCATGCTTTAGAGTTTTGTAAATTTCAAACAAATTAAAATATCTAAAGTAACTATAAAAAAAATAAAATATCTAAAGTAACTATAAAAAAATAAAATATCT
AAAAACCCTTTGAACGAGTTAATATCAAGGAAAAACTAAAATATCTTCAGTGCACTGTTTTCACCAAAAAAATAAAAACATAAAAAACTCCAGTCCACTAGTGTTTTGCACTCA
GTGGCTATGATGACCTATGTAATCATCTATACTAGATAGTAAATAACAAGTTCTACACAAATTGTTCATTTTTTTCTAAAATTTCTTCTAATCTATAGAAGAAAGGTGAAGAATG
TAAAATTTAATATATGTGAAGAATGTGAAAGAGTTCTCACATATATAAAGAAAAAACTCTAACTAACTAAATAACCAATATTTCACTGCAATAGATGGTGGTGGCAGAGCGGA
TATACCTTATGCTTCGACTGCGGTGGTGGAGGACTTGGATAATCAGATTATGGTGGTGTTGTTGTTGAAGGTTTAGCTGCAGTTCCGGAAGGATTTTCACTTCATGATGCTCTT
GACGAGTATGAGTAATTTAATATTGTTGTAGAAGTTGTGGTGAGTTCAACAAGCATTAAGTATACTTTTCCAAATATTTTGGTGAGTTGAATTTTAGATATTTAAAATATAAGTC
CATTTTTATTTATATATCTTATACTATTTGCTTTCTATTTATGCTTGTCTTTTGGAAGTGGAATTGGGGCAACCTACACGTGTCTTACTTTATAGGGTTGATAATGGCATTGAATTG

CTCTGATCCATGTCGATCCAATCGCAGGTATTTTTACTGGCTAAAACTTTTATAGTTTATAGTTAATACTTAATACTCAATGTGATGTCGGCTTGGAGAAGTTTTGTCCTATTCTT
GGAGTAGTGTGACTAGTGTAACTATATACTCTTTCCCAAAATAGATGTTGAAAATAAAAATCACATTTAATAAAATAAAAGTACACAATAATTGAATTATGGTTTTTTGTATTTT
GTTAAAAAAAAATTAAAAAGATTATTGTTAATTATTTATTGACTATGGTGAAAATGAATGAGATTTTATATTAATAATCAACAATTTTTTTAAAAAAAGGTCTTGTATATTTATGT
TATGTTTGGATACACATTGAAAATCATAAAATGCGTGTCTAACATACACTGGTCTTTAAAATGTGTATCCAAAATATGCACTTAGACCATCTCCAATGGGAGTTCTTTGCAAGTT
CTTTAAAGAACCAATTATATATTTTCAATATTTTAATCAATGGAGTGTGCCAAGTTGGAGAACCATTACTTGCTGAAGCAACTTTGGTGATGTGGTTCTCAATAAAATATGATGT
GTCTTAATGGATTTCACAAAAAACTCAAAAATGAGTTGCACCATTGGAGATGCTCTTAGTTTGATGAAGATATATGTTGTTCTTTTTTAATAAATTTGGATAAGAAATATGTGTT
TTTCTTGTTTGTTTATATTTTAGGGCGGAGGTACAAGGTAGTATTATTGGGTCAAAATGTATTTTGGTTCGCGAAGGAAAAAAATTGTATGCTTCATTATCTATTGTAAGTTTAT
GTTTAGTTTCACGTTTAAAATTGATTACGGATGGTTAGAATTGATTTTGACATTTTTATTTGTCTTTGAGTAAAATTGATTTTGATATGTTTGATTGCTATAGAGTTGGATTAATT
ATACTTTTCATAATTAATTGTTCTTGAAGCTAAATTTTTTAACTTTTACGTGTAAACAAGATTTTTATTAAGGGTGACAAAACATATCTCGTTCGCTGGACATGTTTGTTTTGCCCG
CACTATTTCGCAGGATATTCCAAGATTTTAGATCAACACTCTCTAATATGTTGGATTTGCTTCCACTTCGCCATAATTTTTTGTAGGACGAGTCAAAGTTTTAGGCTTGTGCCCTC
AACTATGTCTCTCTCGTCTCACCTTTTTTTTGTGCAGGTTTTTGTGTGGCGAGTCTAAACCGAGCGAGTATGTCCGACACTTAAGCTTAATGTTAACCTCACTTCTACAAAAATAT
ATGCAAACATAAATCAGTTTATCTTTAACTCATTTTTAGAAAGAATCAATTTTATGTGAACTATTCACTCTATCAACTTTCTTTTATGCAAAACCAAACACACATTAAGCATTAAGT
TATATTGATATGTTACTTTATACTTTCATTCATATGCAAATTTAACGTGAAAAGTAAGAAGCTTTAATTTGGAGCATTGGCCAAATGCACAACGGATGGATATCCCATAGTAAAA
TAGAGAAAAAAGATCTTATTATCGGTAGAAAATAGATGCCGAGGGATTTAGAATTCTTAGTTGAATTTGATTTAAAGCTAAACTTCTAAGGTCGTCTCCGACCCACATCTTGAG
ACAGGGTTGCATACTTCGATTTTGTAAGTCTACTTTTGTCATATAATGATAAGATATTTGATGATAATTACTCTAAATGTAGTTCTTTATTCATCTTAATTAACAAATATTTCACAA
TAATGTCTTGTGTTAGAATGTATGATGAGTTTATTTGTCCACTTCTATGAAATCCAATTGATTTTTTAGCAAGTCCTTACTATTTTTAAATGATAAGGAATGCATTTCCTACATATC
GTTGGTTTAAGATGTACAAGAATATTAAAGCACTAAGAGTTTCACCTAAATCTACTTCAATGGTTAAAAATTGAAACTAAGTCCTCGTAGATAAGAGATGAAAAAAAAACATGT
GTTAAAGATACTAATGTACATATAGTTAAAATTCAGTCTCTCTCAATATCACAAACAAAAATTCTATTTGAGGGTGACAAGCTGACATTTCTGCTTTCTCAAAATCTAGGAGTCC
AACCCATTGTGTTCCCTCTAAAACCTATGAGTGACAAACATCTATGTCTACTATATTTAGGCACGAGTTTGTTAAATCCTTTTAAAATGGTCTCATGGACAAAGTAAAGGGGATA
TATTTGATACCTTAGTCTCGTTATGTTAAAAATAGAGATAAATGAGTAGATCAATCTATTTTGTCACTATTTCTTCTTATCTCTTTTTTAAGCATGCATATTAAGAATAGGTAAAA
ATCCTTCTTGATTGTTTTAAATTGAGGTGATTGTCGAAGGTATTTCTTGTTGTAAAAGGGTTCGAGTACCCCTTGTACTCGTTTGAATGTTTATTTAAAAAAAAATATAGGTAAA
GGGAAGTTAAGAAGGGAAAAAAGACATCAAACAAAAATTAAAATCAATGTTTCAGATGATGTAGTCAATTGCTGCCATTTGGGCTAATCAATGAACGTGTCATGAAAATCTTG
GGTTTTGATATATAATGTTGGCTCACTCTAAAATACTTTGTCGATGAAATATTAAATGAATTCAACTGTATCATGCTCGAAAAACTTGTTTTAGACCATATGATTATTGTAGTTTG
AAAACTATGCATTTGTTGCCTATGTCGTTGCTCCCTGCAAATCTCTTATTCGATACTTTATTTAGAAAAGCTATTAATCTGGGCTTATGTTTTTGTATTGACATCATGCTTTAGATT
GTATAAACCAATTTAATGCCAATGTTTCTTACTTCACTTTGCTTAAAAAAAAAAATAAATATGAAATAATTATTACAAAAATCACGATTTCTCATTAAAATTAATAAGATTTTTTTC
TTTGAGAATTATGATTGAAAATTGAACTGTCAACCATATTAAATATTAAACTTTCGTTTATCAACTTATTAATAAAGAAAGATCTAAGTGATACCCTTATCCTAATAATGAAAATT
CACCTTAGATATATCTAAATACTAGATTTTCCAGCTCTCACAAAAGCTTTTGGCATATATATATATATATATATATATATATATATATATATATATATATATATATATATAATATGA
CAAAAATATATTAAACTTAGTAATATGACACCGCCAATAAGTCTTCATCAATTATTTTTATGACAAAATCAATTATTAGATTAAAATCTATTATCATATAAATCATGTCTATAAAA
CTTAATACCAATAAAAATTTATATGATAAATAAAGGGAAAGATCAAATTAACGTTAATAATTATGCATTTTGTTGACGTGTCGAATAATATTTTATTAATGTTCAATTTTATAGAT
ATAATTTATACGATAAAAATTTTAAATTAAATCGTAAATTTTTTTATATGAATAAACGTTAAATCGTAAATTTTTTATTGATGTACGATATTATATATATATATATATATATATATA
TATATATATATATATATATATATATATATATATATATATTCTAGTAATATATCTCTTGACTAAATTATTTTTATTATGATTATAACAGAAATATAAAAGACCACGACCCACCCGACCCA
TACAGAGTTTTCATAGGTTTTTTTTTGACAAGGTCTTTTTTGTTCTTTTGTGTTTGCATTATTTAAATTTTCAAGTATACTTTCACCATATACAGTCTGGACAAAGAGTTTAACATC
CAATAAACACTTGAATTTTATGTATTTTCATCTAAATACAGTAAATTATCATAATTGTCAAATAAAAAAAACATAATAATAATAATAAATGCGAAACCCTGAACTGAAAAATATC

TAGTTCTGAAAGGCGGTTTTCTTGTGCATAGTAGAAGCCGTCACTTACGAAATCACCCCTGCAAGCCACGCACGCTCACCCCACCTCACAACTTGTGCACGATTTTTCCCTTTAA
CCATTAGCCGCTCTATTTTTCATCTCTATCCCAATTTTTCTCATATTCTCCTAAATCAGTAACAGAATCACAAAACCAACTTAAACAAGGACACAGTGAGTTGAGAGTGTTCACAC
TCGAGTCAAAACCCTAACTTACCAGCGATCCGCCATAACTGACGCCGGCGGTGGTTTGACGGTTGTATCCGCGATGGTTGCCGAAAAATCCGTCGTGTGGAAGTGACAATCCG
TTTTGACGGTAGTGATCTCAACGGAACAGAAGGTTGCAGTTCCGGAACAATTTCAACCGAAGGATGTACTTGATCTACACACTTTGCTGGTTTTGATAGTGAGTATGTGTAATT
TCTGGATTTGAGATCCACTACGTGAAGTTCACGCTGTTTGAGTTTTTAAATATTTTGTAACACTGTTTTGTTTTCTTCTTGCAAGTTTTTTGATCCTCTGAAATCCTTGAACCTTCG
TTGGTGAAACTTCATGCTCATTTTTTCTAAACGAAAAAATATATATCTAATATAGCTGTAAGTTTTTTTTTGTTTTCAAAACCGTATTGTATATGCTTTGAATAGGCAATATTTACC
GATGATTTATTTTGAATTGTCCTAAATTAGTTTTAAAGTTATTCTTGGAGTGCAGGGATTATATCATGAGTTATACACTTACACTGTCATCATTCCAGTGTAGGTAAGTTCATGCC
CTTTTATTCTTCTTTTATTTAAACTTTCCTTTTAAGTTTTATA/-

### Variant 3 sequence

TTCTTCTTCTTCTTCTTCTTCTTCTTCTTCTTCTTCTTCTTCTTCTTCTTCTTCTTCTTCTTCTTCTTCTTCTTCTTCTTCTTCTTCTTCTTCTCCTCTAGCCTTTTCA

### Variant 4 sequence

GATTTATGATGTGTTATTAATTTTAATGCTTTATTGTTTTCTGAAAGGCAAATAAGATATGTAATAGATAAAAAGATAAAAACTATTTTACAAACAATCAATAGAAAAATTCAAG
GAATACATAAGAAGTATTAGATCCGAGATCAAAAACGAACATAAAAAACGCAAAACATCTACCAATTACAATAAACTCAGCCGACAATGAACTAGGGCTACCCAATGTGAGG
GAGAACAAAGAAAACACACATAAAAACTTCTTTCTGCTTCAAACGAGACTAGGCTTCTC

### Variant 5 sequence

AATGTTTAATATTTCCAACAAAATTCCCCTGTAAACATTAAATGCTTCTTCT

**Supplementary table 10.** Primers used in the PCR-based marker of the 7441-bp deletion in the *FTa1*-*FTa2* intergenic region.

| Primer name | Sequence |
|---|---|
| F1 | TGGGCTTGATACTTTGTACTCC |
| F2 | TCTACACACTTTGCTGGTTTTG |
| R | CCATCACAATTCAAAGCAATG |

PCR conditions: PCR was performed in a final volume of 40 µL containing 250 ng template DNA, 10 µL of 5x reaction buffer, 10 mM dNTPs, 0.2 µM of each primer, 50 mM MgCl2, 0.2 µL of MangoTaqTM DNA polymerase (Bioline, Australia) and autoclaved Milli-Q water to final volume. Reactions were performed in a thermal using the following program: an initial denaturation of 5 min at 94°C, followed by 40 cycles (94°C for 45 seconds, 58°C for 30 seconds, extension of 45 seconds) and a final extension of 10 minutes at 72oC.

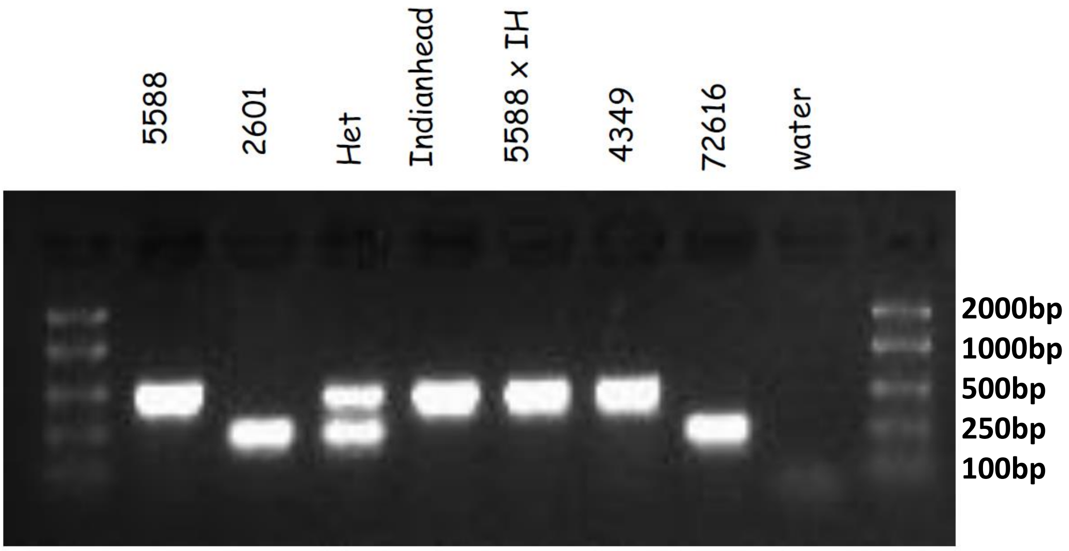


**Supplementary Figure 8.** Visualization, in 2% agarose gel, of the PCR products obtained with primers described in supplementary table 9 tested in different lentil lines. The ~500 bp and ~250 bp bands correspond to primer combination F1/R and F2/R, respectively.

| Accession | species | Country | Latitude | Longitude | Altitude | Accession | species | Country | Latitude | Longitude | Altitude |
|---|---|---|---|---|---|---|---|---|---|---|---|
| ILL 1823 | L. culinaris subsp. culinaris | Afghanistan | 36.9 | 70.9 | 1525 | PI 345631 | L. culinaris subsp. culinaris | Russia | 53.2 | 45 | 220 |
| ILL 2153 | L. culinaris subsp. culinaris | Iraq | 36.8 | 43 | 460 | PI 345552 | L. culinaris subsp. culinaris | Russia | 42 | 47 | 3200 |
| ILL 2214 | L. culinaris subsp. culinaris | Lebanon | 33.8 | 36 | 910 | PI 368649 | L. culinaris subsp. culinaris | Serbia | 45.4 | 19.2 | 620 |
| ILL 223 | L. culinaris subsp. culinaris | Iran | 38.7 | 46.3 | 1360 | PI 374116 | L. culinaris subsp. culinaris | Morocco | 31.8 | - | 480 |
| ILL 2276 | L. culinaris subsp. culinaris | Lebanon | 33.5 | 35.4 | 30 | PI 420924 | L. culinaris subsp. culinaris | Jordan | 32.1 | 35.8 | 860 |
| ILL 2601 | L. culinaris subsp. culinaris | India | 23 | - | - | PI 426775 | L. culinaris subsp. culinaris | Pakistan | 27.6 | 68.1 | 42 |
| ILL 4349 | L. culinaris subsp. culinaris | Canada | - | - | - | PI 426788 | L. culinaris subsp. culinaris | Pakistan | 31.9 | 73.1 | 180 |
| ILL 4370 | L. culinaris subsp. culinaris | Iraq | 33.3 | 44.4 | 40 | PI 426797 | L. culinaris subsp. culinaris | Pakistan | 32.3 | 74.4 | 234 |
| ILL 4605 | L. culinaris subsp. culinaris | Argentina | - | - | - | PI 472291 | L. culinaris subsp. culinaris | India | 26.7 | 85.2 | 50 |
| ILL 5065 | L. culinaris subsp. culinaris | Jordan | 32.6 | 35.7 | 300 | PI 472311 | L. culinaris subsp. culinaris | India | 22.4 | 88.1 | 10 |
| ILL 5588 | L. culinaris subsp. culinaris | Jordan | 32.1 | 35.7 | 700 | PI 472317 | L. culinaris subsp. culinaris | India | 14 | 77 | 600 |
| ILL 5895 | L. culinaris subsp. culinaris | Ethiopia | 7.1 | 39.1 | 2440 | PI 472328 | L. culinaris subsp. culinaris | India | 27.6 | 81.6 | 126 |
| ILL 5976 | L. culinaris subsp. culinaris | Cyprus | 34.8 | 33 | 260 | PI 472343 | L. culinaris subsp. culinaris | India | 26.3 | 78.2 | 180 |
| ILL 6005 | L. culinaris subsp. culinaris | Argentina | - | - | - | PI 472359 | L. culinaris subsp. culinaris | India | 26 | 93 | 60 |
| Indianhead | L. culinaris subsp. culinaris | Canada | - | - | | PI 472578 | L. culinaris subsp. culinaris | Iran | 35.8 | 51 | 1300 |
| PI 297774 | L. culinaris subsp. culinaris | Greece | 40.9 | 23.7 | 140 | PI 472625 | L. culinaris subsp. culinaris | Iran | 35.2 | 59.4 | 1340 |
| PI 297779 | L. culinaris subsp. culinaris | Greece | 39.7 | 20.8 | 580 | PI 509333 | L. culinaris subsp. culinaris | Turkey | 37.2 | 38.8 | 650 |
| PI 297789 | L. culinaris subsp. culinaris | Greece | 38.3 | 20.5 | 700 | PI 509409 | L. culinaris subsp. culinaris | Turkey | 39.7 | 35.8 | 1300 |
| PI 298122 | L. culinaris subsp. culinaris | France | 48.3 | - | 60 | PI 513328 | L. culinaris subsp. culinaris | Pakistan | 25.3 | 68.8 | 20 |
| PI 300248 | L. culinaris subsp. culinaris | Syria | 36 | 37.2 | 400 | PI 533693 | L. culinaris subsp. culinaris | Spain | 42.3 | - | 860 |
| PI 300250 | L. culinaris subsp. culinaris | Syria | 36.4 | 37.2 | 460 | PI 606600 | L. culinaris subsp. culinaris | Nepal | 29.6 | 81.3 | 1368 |
| PI 339281 | L. culinaris subsp. culinaris | Turkey | 38.2 | 29 | 780 | PI 606610 | L. culinaris subsp. culinaris | Tajikistan | 40.4 | 69.9 | 560 |
| PI 339293 | L. culinaris subsp. culinaris | Turkey | 38.3 | 33.6 | 920 | PI 606615 | L. culinaris subsp. culinaris | Russia | 45.8 | 38.6 | - |
| PI 357227 | L. culinaris subsp. culinaris | Macedonia | 42 | 22.2 | 450 | ILWL 7 | L. culinaris subsp. orientalis | Turkey | 38 | - | - |

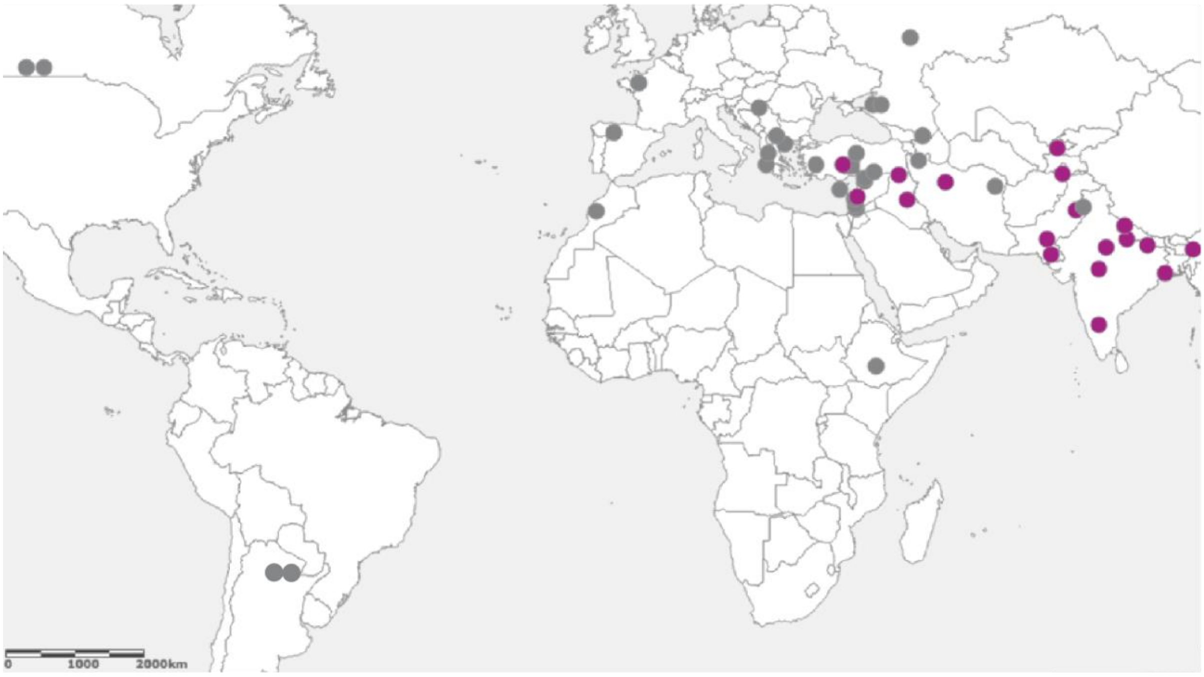


**Supplementary Figure 9.** Incidence of the 7441-bp deletion in 48 lentil lines (University of Tasmania lentil collection) with diverse geographical origin. Purple or grey circles in the map represent accessions that carrying (ILL 2601 allele) or not carrying (ILL 5588 allele) the deletion, respectively.

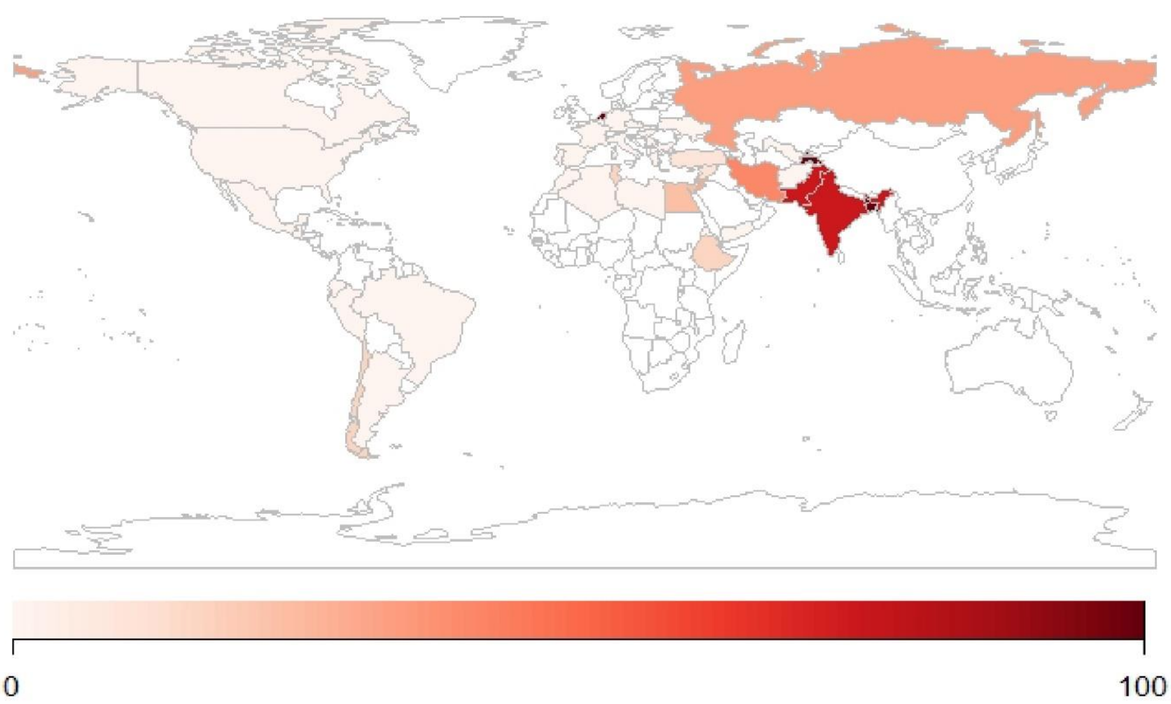


**World**

| | N | WT | DEL | %DEL |
|---|---|---|---|---|
| World | 324 | 249 | 75 | 23.1 |

**Central Asia**

| Country | N | WT | DEL | %DEL |
|---|---|---|---|---|
| Tajikistan | 2 | 0 | 2 | 100.0 |
| Afghanistan | 7 | 7 | 0 | 0 |
| Uzbekistan | 1 | 1 | 0 | 0 |

**South Asia**

| Country | N | WT | DEL | %DEL |
|---|---|---|---|---|
| India | 28 | 7 | 21 | 75.0 |
| Pakistan | 8 | 2 | 6 | 75.0 |
| Bangladesh | 4 | 0 | 4 | 100.0 |

**Africa**

| Country | N | WT | DEL | %DEL |
|---|---|---|---|---|
| Eritrea | 1 | 1 | 0 | 0 |
| Ethiopia | 24 | 20 | 4 | 16.7 |

**America**

| Country | N | WT | DEL | %DEL |
|---|---|---|---|---|
| USA | 6 | 6 | 0 | 0 |
| Peru | 1 | 1 | 0 | 0 |
| Brazil | 6 | 6 | 0 | 0 |
| Argentina | 3 | 3 | 0 | 0 |
| Canada | 28 | 28 | 0 | 0 |
| Chile | 12 | 10 | 2 | 16.7 |
| Guatemala | 2 | 2 | 0 | 0 |
| Ecuador | 1 | 1 | 0 | 0 |
| Mexico | 6 | 6 | 0 | 0 |

**Middle east**

| Country | N | WT | DEL | %DEL |
|---|---|---|---|---|
| Jordan | 10 | 7 | 3 | 30.0 |
| Yemen | 2 | 2 | 0 | 0 |
| Syria | 11 | 10 | 1 | 9.1 |
| Iran | 49 | 29 | 20 | 40.8 |

**Mediterranean**

| Country | N | WT | DEL | %DEL |
|---|---|---|---|---|
| Greece | 5 | 5 | 0 | 0 |
| Egypt | 13 | 10 | 3 | 23.1 |
| Morocco | 5 | 5 | 0 | 0 |
| Spain | 4 | 4 | 0 | 0 |
| Tunisia | 6 | 5 | 1 | 16.7 |
| Algeria | 2 | 2 | 0 | 0 |
| Libya | 4 | 4 | 0 | 0 |
| Portugal | 2 | 2 | 0 | 0 |
| Turkey | 20 | 18 | 2 | 10.0 |
| Lebanon | 3 | 3 | 0 | 0 |
| Israel | 2 | 2 | 0 | 0 |
| Italy | 3 | 3 | 0 | 0 |
| Bulgaria | 1 | 1 | 0 | 0 |
| Macedonia | 4 | 4 | 0 | 0 |
| Serbia | 1 | 1 | 0 | 0 |

**Europe**

| Country | N | WT | DEL | %DEL |
|---|---|---|---|---|
| Russia | 3 | 2 | 1 | 33.3 |
| Ukraine | 1 | 1 | 0 | 0 |
| Belgium | 1 | 1 | 0 | 0 |
| Netherlands | 1 | 0 | 1 | 100.0 |
| Czech Republic | 4 | 4 | 0 | 0 |
| Hungary | 3 | 3 | 0 | 0 |
| France | 4 | 4 | 0 | 0 |
| Germany | 2 | 2 | 0 | 0 |

**Other**

| Country | N | WT | DEL | %DEL |
|---|---|---|---|---|
| Unknown | 2 | 2 | 0 | 0 |
| ICARDA | 11 | 7 | 4 | 36.4 |
| USDA | 5 | 5 | 0 | 0 |

**Supplementary Figure 10.** Incidence of the 7441-bp deletion in a panel of 324 lentil lines with diverse geographical origin. Colour in the map represent percentage of accessions that present the deletion. Countries without colour were not represented in the panel. N = total number of accessions; WT = number of accessions with a wild type allele (no deletion present); DEL = number of accessions with the 2601 allele (deletion present); %DEL = percentage of accessions carrying the deletion.
